# Computational design of N-linked glycans for high throughput epitope profiling

**DOI:** 10.1101/2023.04.04.535514

**Authors:** Per Greisen, Li Yi, Rong Zhou, Jian Zhou, Eva Johansson, Tiantang Dong, Haimo Liu, Laust B Johnsen, Søren Lund, L. Anders Svensson, Haisun Zhu, Nidhin Thomas, Zhiru Yang, Henrik Østergaard

**Affiliations:** Data Science and Innovation; Global Research Technologies; Discovery Technology China; Global Drug Discovery, Novo Nordisk A/S

**Author notes:** Corresponding authors: Per Jr. Greisen Digital Science and Innovation, Novo Nordisk A/S Rong Zhou Discovery Technology China, Novo Nordisk A/S Henrik Østergaard Global Drug Discovery, Novo Nordisk A/S.

## Abstract

Efficient identification of epitopes is crucial for drug discovery and design as it enables the selection of optimal epitopes, expansion of lead antibody diversity, and verification of binding interface. Although high resolution low throughput methods like X-ray crystallography can determine epitopes or protein-protein interactions accurately, they are time-consuming and can only be applied to a limited number of complexes. To overcome these limitations, we have developed a rapid computational method that incorporates N-linked glycans to mask epitopes or protein interaction surfaces, thereby providing an atomistic mapping of these regions. Using human coagulation factor IXa (fIXa) as a model system, we could rapidly and reliably delineate epitopes through the insertion of N-linked glycans that efficiently disrupt binding in a site-selective manner. To validate the efficacy of our method, we conducted ELISA experiments and high-throughput yeast surface display assays. Furthermore, X-ray crystallography was employed to verify the results, thereby recapitulating through the method of N-linked glycans an atomistic resolution mapping of the epitope.

## Introduction

The ability to rapidly identify epitopes and protein-protein interactions (PPI) is of utmost importance not only in interrogating new biology (1) but also in drug design, where identification and selection of optimal epitopes represent a key element in the optimization of potency and safety (2). It is important to have geometrically optimal epitopes to achieve the optimal potency of multispecific antibodies. This is particularly important in the case of bispecific antibodies like Mim8 (3) and emicizumab (4). Moreover, in *de novo* computational protein design, it is essential to rapidly determine whether the design is binding to the desired epitope. Here, the binding is often tested through methods like yeast surface display (YSD). Hence, there is a need for a fast method to structurally pinpoint the position of the epitope or determine if the binding interface is in accordance with computational design.

Traditional methods for mapping epitopes have certain limitations associated with them. One commonly used method is binning monoclonal antibodies (mAbs) into different epitope bins, thereby grouping the mAbs into overlapping or similar epitopes (5). Using full-length mAbs can be misleading due to steric hindrance of the fragment crystallizable (Fc) region, which does not necessarily reflect the binding region of the epitope. Other common ways of determining epitopes include mutational scanning, peptide mapping, NMR spectroscopy (6, 7), X-ray crystallography (8, 9), and cryo-electron microscopy (10). However, these are considered low throughput methods that require considerable effort or time and can test only a limited number of complexes. The first two methods have low resolution, while X-ray and cryo-EM of the co-complex can provide atomistic details of the interaction. Another popular method that can be used to profile epitopes and protein-protein interactions is hydrogen-deuterium exchange mass spectrometry (HDX-MS), which has been used to accurately map interactions between antibodies and antigens (11). The technique can accurately identify epitopes, but it is important to be cautious when analyzing the results as significant changes upon binding of the overall structure of the molecule can make it hard to interpret. In proteins with allosteric sites it can be difficult to map the precise epitope using HDX-MS due to conformational changes upon binding (12).

N-linked glycosylation is a common post-translational modification that occurs in many human proteins (13). It expands the functional properties of proteins by increasing the physico-chemical properties, e.g., stability (14). N-linked glycosylation of proteins occurs through a consensus Asn-Xxx-Ser/Thr recognition motif. Here, the sidechain of the initial Asn represents the glycan acceptor site; the second position (Xxx) can be any amino acid except proline and is followed by either serine(S) or threonine(T) at the third position. The efficiency of a glycosylated site depends on many factors, including the local 3-dimensional structure around the consensus motif (15) and surface accessibility. Introducing a N-linked glycan within an epitope will disrupt ligand binding through simple steric hindrance. Others have investigated the idea of using a N-linked glycan to mask or protect an epitope with great success. Lombana et al. (16) developed the glycosylation-engineered epitope mapping (GEM) method, which involves the insertion of N-linked glycans at specific sites on the surface of the antigen to conceal localized regions. The method calculates the solvent accessible surface area (SASA) of the N and S/T residues and determines if they are exposed. There is no structural modeling of the N-linked glycan, and the method has not incorporated design of the S/T residue at the third position, which can have different preferences depending on local protein structure.

As discussed, identifying epitopes in a high throughput manner with high precision is desirable to understand biology and engineer optimal drugs targeting specific epitopes. Hence, we have improved on existing computational methods to insert and design the local sequence of the target protein such that it can accommodate a N-linked glycan at scale. It was achieved through a combination of structural modeling and computational design techniques and can be fully automated. Further, it is possible to add N-linked glycans at multiple positions of the human factor IXa without altering its overall structure (17). Using this model system of factor IXa, we were able to validate the method in accordance with a crystal structure of the complex using low-and high-throughput experimental methods.

## Methods

### Generation of anti-FIXa antibodies

Antibodies specific for human FIXa were generated in Kymouse mice (Kymab Group Ltd) that comprise a human antibody repertoire. Kymouse mice were immunized with active-site inhibited FIXa purchased from Haematologic technologies Inc/Prolytics. After electrofusion, hybridomas were plated in microtiter plates and supernatants were screened by a FIXa ELISA. Hybridomas secreting antibodies specific for FIXa were propagated and their supernatants subsequently used for proteinA affinity purification. FIXa binding was confirmed on fortebio, and hybridoma hits were sequenced and recombinantly expressed in HEK293 cells using standard techniques.

### N-linked glycan library production and characterization

The DNA fragment encoding the amino acid sequence of HPC4 tagged human factor IX (fIX) without the gamma-carboxyglutamic (GLA) domain was synthesized and cloned into the pJSV vector with CD33 signal peptide to replace the native signal peptide(18). 98 variants were created, each containing specific engineered mutations for N-linked glycosylation. The mutations introduced were an asparagine (N) residue followed by any amino acid residue except proline (X) and a serine (S) or threonine (T) residue at different positions in the DNA sequence. Each of the 98 variants were individually cloned for further study or experimentation. HEK293 6E cells were used for transient production of these proteins following standard protocols for 293Fectin (Invitrogen). The cell cultures were harvested at 5 d post-transfection by centrifugation at 6,000 rpm for 15 min, and the supernatants were filtered using a 0.45μm filter for purification with anti-HPC4 sepharose 4FF affinity columns ( anti-HPC4 antibody coupled to the CNBr activated sepharose 4FF resin, GE), equilibrated in buffer (20mM Tris-HCl, 100mM NaCl, 1mM CaCl_2_, pH 7.4). The bound proteins were eluted with buffer (20mM Tris-HCl, 100mM NaCl, pH 7.4, 1mM EGTA) as final products. The variants were analyzed with SEC-UPLC, LC-MS and SDS-PAGE, which were also digested with human factor XIa(19, 20) and then analyzed with SDS-PAGE to identify if the N-linked glycosylation was inserted.

### FACS analysis

The yeast surface display plasmids were constructed to evaluate the binding of single-chain variable fragments (scFv) to different factor IXa variants. The *AGA2* gene downstream of the *GAL1* promoter was fused with a four-part cassette encoding the VL, linker, VH, and FLAG tag fragments, forming an VL–Linker–VH-FLAG-Aga2 cassette. Two plasmids that contain (G4S)3 and (G4S)4 linker, respectively, were constructed.

The constructed plasmids were transformed into EBY100 cells (*URA*^+^, *leu*^−^, *trp*^−^) followed by cultivation and induction. The cells bearing different constructs were labeled with anti-FLAG-iFluor 488 and anti-HPC4-iFluor 647 antibodies (GenScript, China) followed by detection using similar protocols published previously(21) with the NovoCyte flow cytometer (Agilent, USA). The iFluor 647 fluorescent intensity was detected with the APC channel, 670/30 nm band pass, and iFluor 488 fluorescent intensity was detected with the FITC channel, 530/30 nm band pass. The binding efficiencies against different fIXa variants were evaluated as: Binding efficiency = [Mean Fluorescency Intensity of iFluor 647] /[ Mean Fluorescency Intensity of iFluor 488], in which Mean Fluorescency Intensity (M.F.I.) of iFluor 647 and M.F.I. of iFluor 488 represent the binding of fIXa variants and surface display of scFv, respectively.

### ELISA of N-linked glycosylation library screening

An anti-HPC4 monoclonal mouse antibody was coated onto 96-well ELISA plates (9018, Corning) at 5 ug/ml, 100 ul/well. The plates were incubated at 4℃ overnight. The plates were washed 3 times with PBS, blocked with 1% BSA for 3 hours. The plates were washed 3 times before adding human factor IX or human factor IX with N-linked glycosylation at 1ug/ml, 100 ul/well. 1 hours later, the plates were washed 3 times before adding anti-FIX antibodies to be assessed at 1ug/ml, 100 ul/well. The plates were washed 3 times again before adding anti-hIgG-HRP (Invitrogen, Cat # A18823) antibody for detection. 30 mins later the plates were washed followed by measuring ELISA signaling at an absorbance of OD450.

### Computational design of N-linked glycans

The crystal structure of fIXa (PDB ID: 1RFN(22)) was used as a starting point for calculating the Define Secondary Structure of Proteins (DSSP)(23) using RosettaScripts (24, 25). A glycan model was generated by utilizing the crystal structure (PDB ID: 4Z7Q(26)) with the glycan alpha-D-mannopyranose-(1–3)-[alpha-D-mannopyranose-(1–6)]beta-D-mannopyranose-(1–4)-2-acetamido-2-deoxy-beta-D-glucopyranose-(1–4)-2-acetamido-2-deoxy-beta-D-glucopyranose (chain K in crystal structure) creating the parameter file used in Rosetta. Protein structures were analyzed and visualized using PyMOL(27). For detail and list of commands and scripts used please see Appendix.

### Production of NN-2003Fab

The DNA fragments encoding the amino acid sequence of fab (NN-2003Fab) heavy chain and light chain were synthesized and cloned into the pJSV vector (generated in-house using a *CMV* promoter) with CD33 signal peptide to replace the native signal peptide, respectively, which then were co-transfected into HEK293 6E cells for expression together following standard protocols for 293Fectin (Invitrogen). The cell cultures were harvested at 5 d post-transfection. Cells were removed from the culture by centrifugation at 6,000 rpm for 15 min, and the supernatants were filtered using a 0.45μm filter. The supernatants were loaded to HiTrap Protein G HP (5ml, GE), equilibrated in PBS. The bound proteins were eluted with elution buffer (0.1 M glycine-HCl, pH 2.7), the eluate was neutralized with buffer (1M Tris, pH9.0), which was further polished with HiLoad Superdex 200 16/60 (GE), equilibrated and eluted with buffer (25mM HEPES, 150mM NaCl, pH7.4).

### Crystal structure determination of NN-2003Fab in complex with des-(Gla-EGF1) fIXa

Crystals of NN-2003 Fab mixed in a 1:1 molar ratio with human EGR-chloromethylketone active-site inhibited des-(Gla-EGF1) fIXa from Cambridge ProteinWorks were grown using the sitting drop vapor diffusion technique at 18 °C. A protein solution of 100 nl 6.9 mg/ml complex in 20mM Tris-HCl, pH 7.4, 50mM NaCl, 2.5mM CaCl2 was mixed with 100 nl of 0.5 M ammonium sulphate, 0.1 M sodium citrate pH 5.6, 1.0 M lithium sulphate as precipitant and incubated over 60 µl precipitant. The crystal was cryo protected in a solution consisting of 0.375 M ammonium sulphate, 75 mM sodium citrate pH 5.6, 0.75 M lithium sulphate, 4 % glycerol, 4 % ethylene glycol, 4.5 % sucrose, and 1 % glucose prior to flash cooling in liquid nitrogen. Diffraction data were collected at 100K at the Swiss Light Source beamline X06DA (1.00000 Å wavelength) using a Pilatus2M pixel detector from Dectris. Autoindexing, integration and scaling of the data were performed with programs from the XDS package(28). The asymmetric unit contains one Fab:FIXa complexes as judged from Matthew’s coefficient analysis. The structure was determined by molecular replacement with Phaser(29) as implemented in the program suite Phenix(30) with the crystal structure of a homologous Fab and FIXa from PDB entry 3KCG(31) as search models. The correct amino acid sequence for the Fab was model built using COOT(32) and thereafter the structure was refined using steps of Phenix refinement(33) and manual rebuilding in COOT. Diffraction data and refinement statistics are found in Supplementary Table 1.

## Results

### Computation-Guided epitope mapping of factor IXa

To profile the epitopes recognized by a panel of antibodies raised against fIXa with high throughput and atomistic resolution, an N-linked glycan variant library of fIXa was generated using a computational workflow (see Figure 1A). The computational protocol was generated to design N-linked glycans for high-throughput epitope profiling with atomistic resolution (see Figure 1B). Firstly, the crystal structure of fIXa, along with a model glycan was used as the input. The glycan was structurally inserted and modeled using RosettaMatch (34) on positions that were considered loops according to the DSSP algorithm in RosettaScripts (24, 25), resulting in 158 positions. From a statistical analysis, an epitope is comprised on average of 9–22 amino acid residues (35), and loop regions are often the most frequent regions used to insert N-linked glycosylation in human proteins. From the computational screen, number of positions were identified, all of which were designed to obtain the N-linked glycosylation motif N-X-S/T (where X represents all amino acids except proline). To increase the quality of the designed N-linked glycans, the 3-dimensional structures were ranked using RosettaScripts to compute the cutoffs on the interface energies, < 0 REU, and geometrical restraints, < 10, based on the structural models. This resulted in 98 positions that, according to the computational design, could accommodate N-linked glycosylation and were chosen for experimental verification (see Supplementary Table 3 for a full list of sequences used in the study).

**Figure 1.**
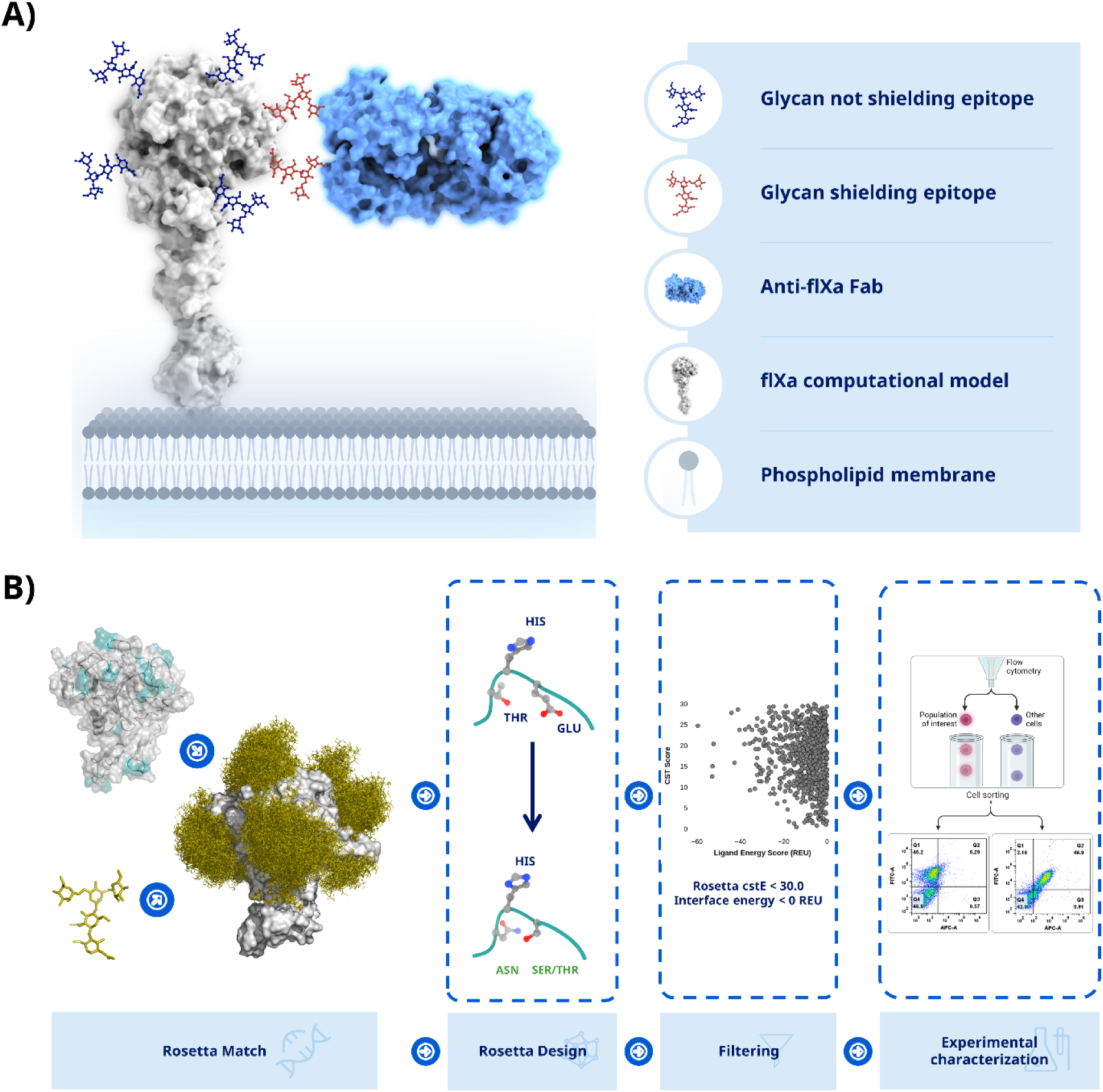
Epitope mapping through engineered N-linked glycans. A) The idea is to engineer N-linked glycans into the protein of interest which will mask the epitope/interface and thereby prevent binding/interaction. B) Firstly, loop elements (cyan) are identified using DSSP on a crystal structure of factor IXa which returns potential positions to place the N-linked glycan. Next, a model of glycan structure is generated which is used by RosettaMatch to insert the N-linked glycan onto the structure of fIXa. B) The sequence is designed by RosettaDesign to obtain the consensus motif N-X-S/T (X can be any amino acid except proline) and optimized according to the input geometry. C) Engineered variants are filtered based on deviations from the specified geometry, < 10, and an interface energy < 0 REU is required. D) The remaining designs are used to experimentally characterize binding epitopes using either yeast surface display or other binding methods such as ELISA.

It was decided to use fIXa without the gamma-carboxyglutamic acid-rich (GLA) domain, as the epitopes of interest were assumed to be distal to the membrane. Ninety-nine variants, including the wild type (WT), were expressed in the mammalian cell line HEK293 to ensure that they were glycosylated. It was possible to purify 92 of the engineered variants which was done in 2 steps. The first batch was produced in 30 ml cell culture while for variants not expressing here were upscaled to 300 mL cell culture. This resulted in concentrations ranging of 0.08 to 2.25 mg/mL for the 92 variants which were used for further characterization. To determine whether the engineered variants had successfully incorporated an N-linked glycan, the level of N-glycosylation of each variant was estimated by separating them based on their mass differences using SDS-PAGE. This method was used to detect and compare the molecular weights of the different protein variants, confirming whether the N-linked glycan had been successfully introduced. Ninety-eight desGla-hfIX variants were designed with a single *de novo* N-linked glycan site, with 77 variants in the protease domain, 12 variants in the EGF2 domain, and 9 variants in the hinge region (see Supplementary Figure 1). SDS-PAGE analysis was suitable for characterizing the glycosylation level only for variants in the protease domain. SDS-PAGE analysis showed that 52 out of the 77 variants were fully or partially glycosylated (for a comprehensive description, see Supplementary Table 2). Since the mass difference of the EGF domains was too small to be detected using SDS-PAGE, the EGF2 variants that were engineered with N-linked glycosylation were tested using LC-MS. Seven of the variants in the EGF2 domain were shown to have incorporated N-linked glycosylation (see Figure 2). Table 1 shows the range of the incorporation efficiencies from 0 to 100% where five of the engineered variants were unquantifiable.

**Figure 2.**
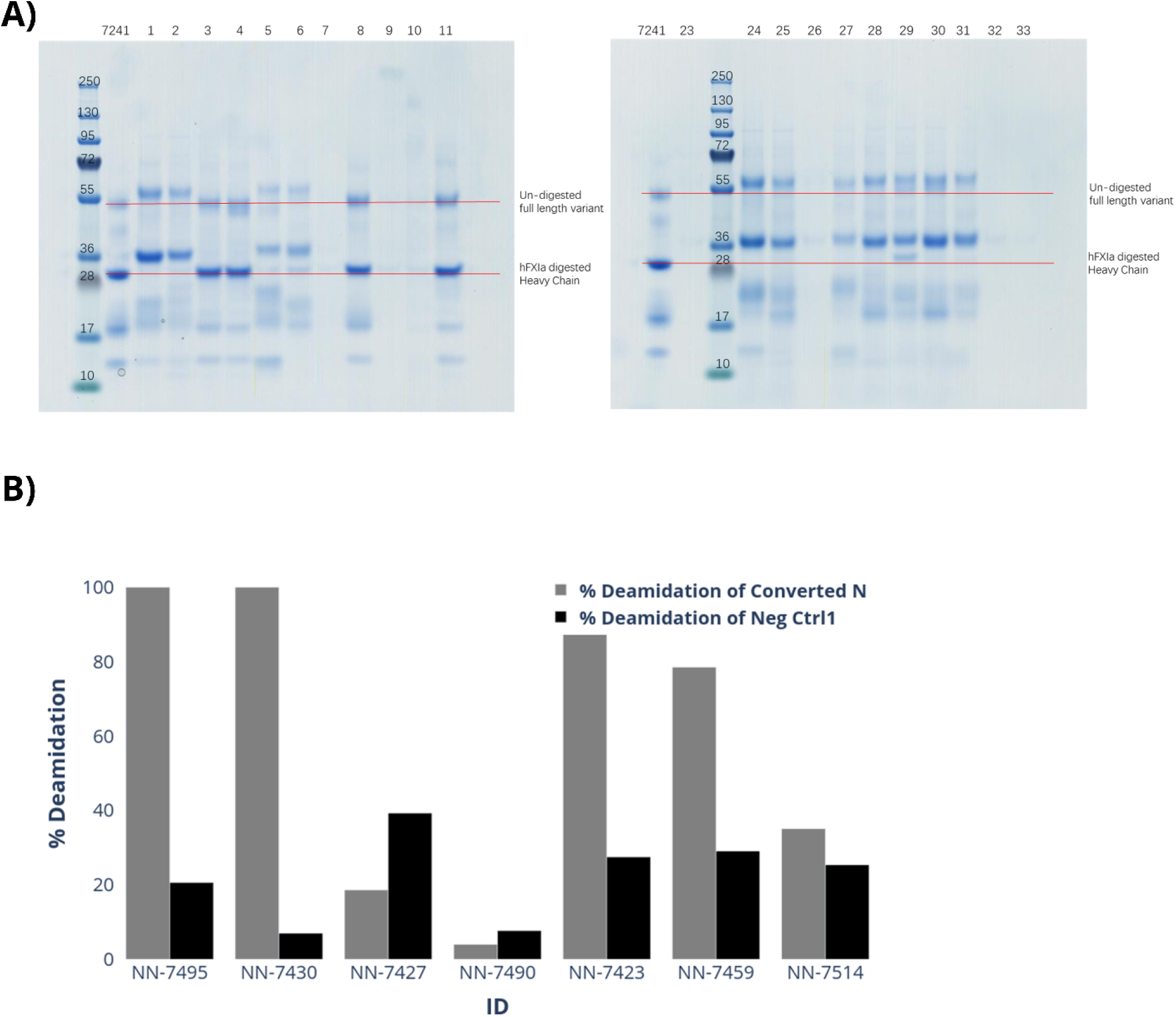
Characterization of N-linked glycosylation in EGF2 domain. A) The engineered variants are characterized on a reducing SDS-PAGE gel with NN-7241 (fIXa without gamma-carboxyglutamic acid domains) as reference. Shifts are indicated with respect to the heavy chain of fIXa (red line) B) The glycosylation level of engineered fIXa variants could be characterized by LC-MS/MS according to convert Asparagine to Aspartic Acid with removing N-linked glycosylation. To mitigate the false positive effect induced from chemical deamidation, deamidation ratio of N175 was calculated as a negative control. As an example, the LC-MS/MS analysis indicated that 3 of 7 variants were 100% incorporated with N-linked glycosylation; 1 variant was partially (78.5% or lower) incorporated with N-linked glycosylation and 3 variants was not or limited (35.1∼4.0% or lower) incorporated with N-linked glycosylation.

**Table 1.** Quantification of the percentage of N-glycosylation presenting in EGF2 variants with peptide mapping.

| **NNCD** | **Serial Number** | **Glycosylation (%)** | **Mutations** | **Position** |
| --- | --- | --- | --- | --- |
| NN-7501 | 82 | 0 | T66N G68S | EGF2 |
| NN-7437 | 18 | 27 | A72N N74S | EGF2 |
| NN-7470 | 51 | 56.8 | S56N D58S | EGF2 |
| NN-7493 | 74 | 57.3 | K60N V62S | EGF2 |
| NN-7435 | 16 | 67.2 | Q75N | EGF2 |
| NN-7438 | 19 | 100 | E73N Q75T | EGF2 |
| NN-7460 | 41 | 100 | A57N N59S | EGF2 |
| NN-7427 | 8 | ND | T41N N43S | EGF2 |
| NN-7430 | 11 | ND | D58N K60T | EGF2 |
| NN-7456 | 37 | ND | E67N Y69S | EGF2 |
| NN-7469 | 50 | ND | E50N F52S | EGF2 |
| NN-7490 | 71 | ND | S77N E79S | EGF2 |

ND means “not determined”.

### ELISA assay mapping of factor IXa epitope

To map the epitope of the anti-fIXa NN-2003 full length monoclonal antibody (mAb), 56 purified variants were first tested in an ELISA assay along with the WT fIXa. Out of these, four N-linked glycan variants (see Table 2) showed reduced signals in the binding assay comparable to the negative control buffer (see Figure 3A). This indicates that the N-linked glycan in these variants interfered with binding and is part of the epitope. Loss of binding was defined as an ELISA signal equivalent to that of the buffer used. To verify if this was due to site-selective masking of the epitope on fIXa, a crystal structure of the co-complex was solved. From the co-complex structure, it was observed that the inserted N-linked glycans were indeed in the binding epitope (see Figure 3B), thus validating the reduced binding signal observed by the ELISA.

**Figure 3.**
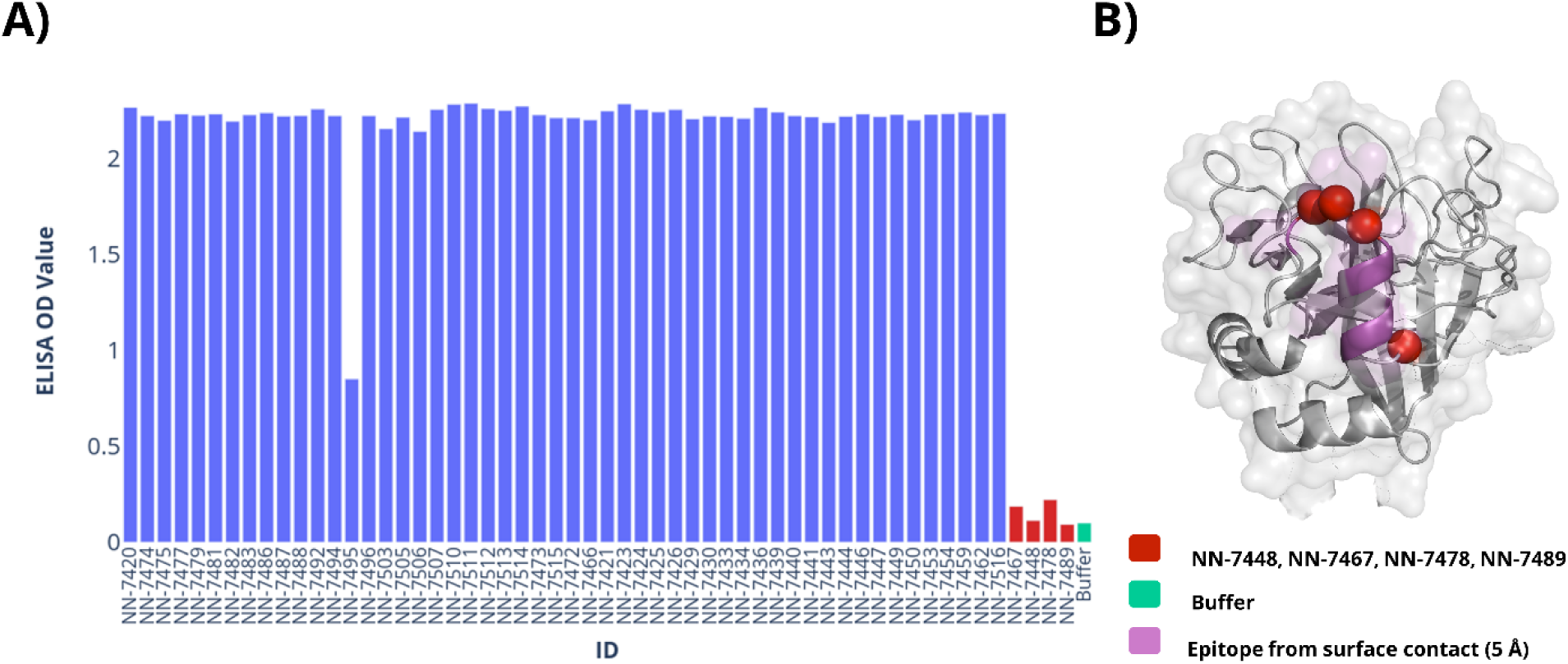
ELISA mapping of epitope. A) Measuring ELISA signaling for each of the N-linked glycosylated variants displaying the maximum value. Variants not showing reduced signal are in blue and the signal from the buffer (green) while variants with signal similar to the buffer are considered none binding (red). B) The 4 variants with reduced signal and their positions (red spheres for C-alpha position) relative to the epitope as defined by the crystal structure (purple).

**Table 2.** N-linked glycosylated variants with reduced binding to NN-2003.

| **NNCD** | **Mutations** | **Epitope** |
| --- | --- | --- |
| **NN-7448** | F296N I298S | Yes |
| **NN-7467** | V285N R287T | Yes |
| **NN-7478** | T294N F296S | Yes |
| **NN-7489** | K295N | Yes |

### Epitope Mapping Using Yeast Display

The first step in raising binders or verifying *de novo*-designed proteins often involves relying on display technology, such as phage panning or yeast display. Compared to phage display, yeast surface display (YSD) can provide a more straightforward quantitative evaluation. In this study, we used two scFv versions of NN-2003 with different linkers, G4Sx3 and G4Sx4 (NN-2003-L1 and NN-2003-L2, respectively), for epitope profiling using YSD. Four N-linked glycosylated fIXa variants were chosen to evaluate the sensitivity of the method, with three N-linked variants in the epitope and one outside of the epitope. Out of the four variants that exhibited reduced binding signals, the three variants with N-linked glycans in the epitope exhibited a reduced binding signal that was comparable to background levels. However, in the case of NN-7505, a binding signal was still observed in a concentration-dependent manner using fIXa as bait. This is in accordance with the binding mode and epitope observed in the crystal structure between NN-2003 and fIXa (see Figure 4).

**Figure 4.**
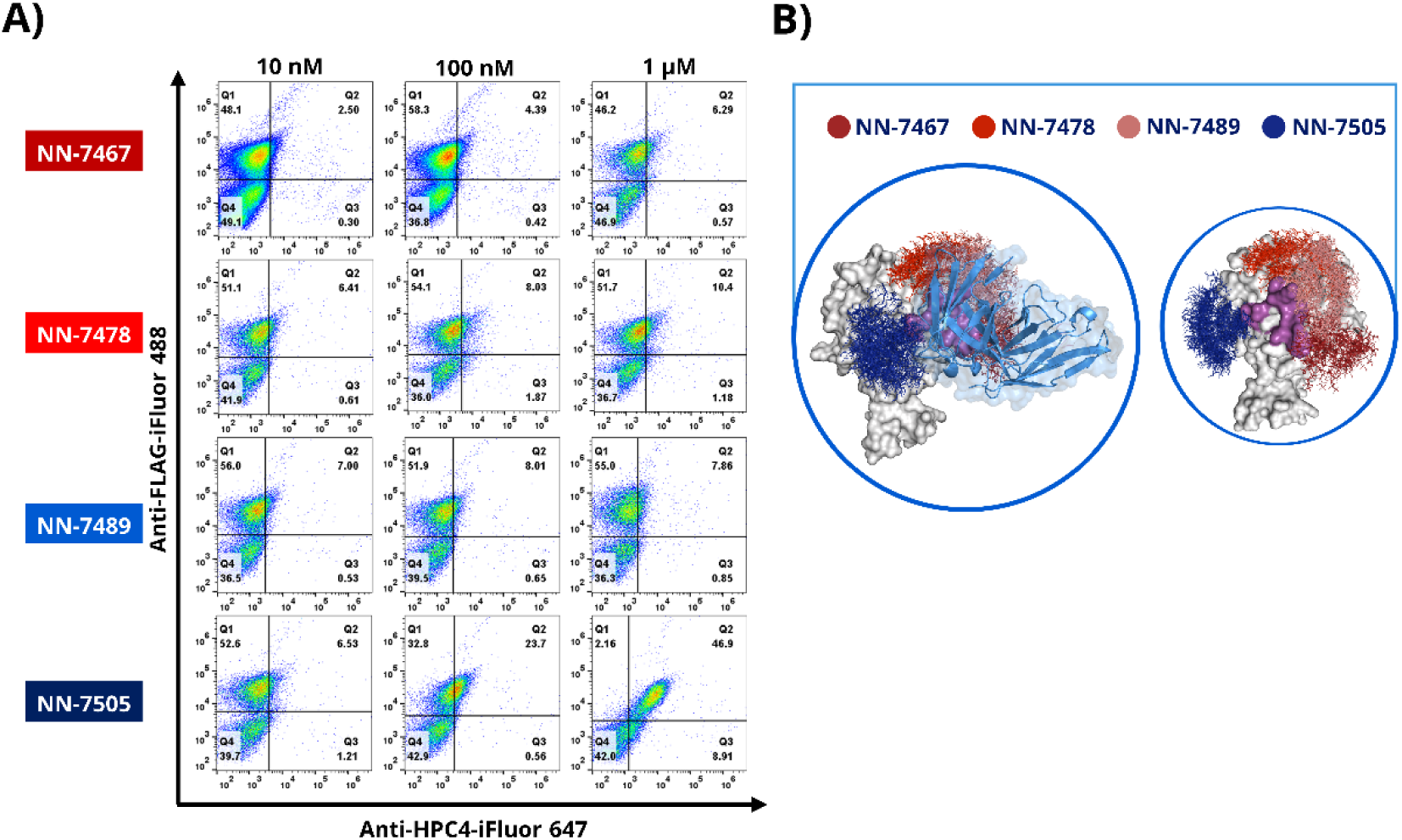
Epitope mapping by yeast display. A) Yeast surface displayed scFv NN-2003 (FLAG coupled with anti-FLAG-iFluor 488, y-axis) binding of 4 fIXa variants with N-linked glycans (coupled with anti-HPC4-iFluor 647, x-axis) on protein titrated at three different concentrations: 10 nM, 100 nM, and 1 uM. B) Crystal structure of the complex between NN-2003’s fragment antigen-binding region (skyblue cartoon representation) and fIXa (white surface repsentation) with N-linked glycans modeled onto the structure where red colors represent epitopes being masked by N-linked glycan while blue indicates no or little effect on binding.

## Discussion

Rapid identification and characterization of epitopes are of utmost importance to optimizing drugs so that they can interact optimally with their targets. Rapidly profiling epitopes with high resolution can, for example, aid in drug discovery to also ensure that lead compounds have diverse interactions with their target, and lastly, validate *de novo*–designed proteins. The ability to rapidly and accurately map epitopes on protein structures is essential for accelerating the engineering and design of proteins, giving confidence in the epitope selection. Here, we have created a fast, reliable, and fine-grained method to insert N-linked glycans using *in silico* methods that map them directly onto a crystal structure or a structural model of the target protein with the necessary sequence modifications to incorporate the N-linked glycan. We used a model system that applies the human coagulation fIXa, which was chosen to verify the protocol. It was shown that the method was robust and could be applied in both low-and high-throughput screening assays, such as ELISA and YSD. In the process of inserting the N-linked glycans, a conservative approach was chosen to mainly focus on loops as defined computationally by DSSP. It has been shown that loops have a higher probability of N-linked glycosylation in human proteins (26) but N-linked glycans are also observed in other secondary elements such as sheets and helices. This method can be easily extended to other secondary elements like helices or sheets by modifying the computational method to include these elements. In this study, to achieve full coverage of the fIXa surface, N-linked glycans were introduced with a maximum spacing of 19 residues between each glycan. As this distance has been observed in other studies of epitopes to ensure coverage of the area, it ensures full coverage of the target protein and increases the likelihood of masking all potential epitopes, even on large proteins like fIXa. Linear epitopes can be in the range of 4-12 amino acids(36) and one might need to expand the frequency of N-linked glycans to other secondary elements in the protein of interest. We did observe that not all the engineered N-linked glycans were incorporated into fIXa. Not all designed positions can sustain the incorporation of an N-linked glycan, and this needs to be considered when applying the method. In a few cases, the site was also not 100% glycosylated, which could potentially make interpretation difficult if the glycosylation level was not properly analyzed. The N-linked glycosylation-dependent method is constrained by the requirement that the target proteins must be expressed in host systems that can perform N-linked glycosylation. Therefore, proteins expressed only in *Escherichia coli* are unsuitable for this method. However, this computational approach is adaptable to any chemical modifications, such as PEGylation, which may offer greater flexibility in some cases. The method’s scope is generalizable to these modifications, making it a versatile technique.

In the ELISA experiment, variability of the signal was observed, indicating heterogeneity for some of the variants. The heterogeneity observed in the binding signal can also be attributed to the occupancy of N-linked glycans, which may result in a reduced binding signal, although this does not necessarily mean the complete removal of the signal. This means that even if an N-linked glycan is present on the protein surface and partially masking the epitope, the scFv may still be able to bind to the protein with reduced affinity. However, we were able to get 59/92 N-linked glycosylated variants with N-linked glycans inserted into fIXa. Using ELISA from the binding results, we were able to determine the right epitope according to the crystal structure with a clear signal of where the epitope was located. Four of the N-linked glycan variants showed a reduced signal, and according to the crystal structure, they were the only variants that should disrupt binding. The computational de-novo design of protein binders is often evaluated using yeast display, but confirmation of epitope binding can be a time limitation. It was not clear whether applying the epitope mapping using YSD would be as sensitive as the ELISA. However, the results showed that we were able to map the epitope using YSD by using four variants with engineered N-linked glycans close to the epitope. Three of the N-linked glycans showed reduced signals in ELISA, and a similar result was observed using titrations in YSD. The results were encouraging, as there were multiple nonspecific interactions between the engineering protein and its N-linked glycans that could be anticipated. To conclude, we have developed an *in silico* method to insert N-linked glycans using structural modeling. The method was validated using fIXa as a model system, and successfully performed epitope mapping using experimental methods, such as ELISA and yeast surface display.

## Tables

**Supplementary Table 1.** *Diffraction data and refinement statistics for the crystal structure determination of NN-2003Fab in complex with des-(Gla-EGF1) fIXa*.

|  |  |
| --- | --- |
| **Wavelength (Å)** | 1.0000 |
| **Resolution range (Å)** | 49.98-2.602 (2.695-2.602) |
| **Space group** | I 2 2 2 |
| **Unit cell (Å, deg)** | 61.44 97.27 366.47 90 90 90 |
| **Total reflections** | 160379 (8632) |
| **Unique reflections** | 33502 (2497) |
| **Multiplicity** | 4.8 (3.5) |
| **Completeness (%)** | 96.24 (73.60) |
| **Mean I/sigma(I)** | 10.62 (1.28) |
| **Wilson B-factor (Å<sup>2</sup>)** | 50.02 |
| **R-merge** | 0.1322 (0.806) |
| **R-meas** | 0.148 (0.9306) |
| **R-pim** | 0.06544 (0.452) |
| **CC1/2** | 0.996 (0.69) |
| **CC\*** | 0.999 (0.904) |
| **Reflections used in refinement** | 33141 (2487) |
| **Reflections used for R-free** | 1657 (124) |
| **R-work** | 0.2670 (0.3629) |
| **R-free** | 0.3236 (0.4278) |
| **CC(work)** | 0.870 (0.672) |
| **CC(free)** | 0.828 (0.538) |
| **Number of non-hydrogen atoms** | 5672 |
| **macromolecules** | 5615 |
| **ligands** | 20 |
| **solvent** | 37 |
| **Protein residues** | 725 |
| **RMS(bonds)** | 0.003 |
| **RMS(angles)** | 0.56 |
| **Ramachandran favored (%)** | 93.28 |
| **Ramachandran allowed (%)** | 6.44 |
| **Ramachandran outliers (%)** | 0.28 |
| **Rotamer outliers (%)** | 3.22 |
| **Clashscore** | 5.76 |
| **Average B-factor ($\text{\AA}^2$)** | 61.15 |
| **macromolecules** | 61.22 |
| **ligands** | 74.69 |
| **solvent** | 43.23 |
| **Number of TLS groups** | 1 |
| **PDB entry** | 8OL9 |

Statistics for the highest-resolution shell are shown in parentheses.

## Acknowledgement

The authors thank the entire Novo Nordisk Mim8 project team for contributions, in particular, Xiaoli Yan for expression of these variants in the library, Xinping Qu, Lin Li, Chenxi Shen for purification of these variants, Xiaoai Wu for collecting data. We would like to express our gratitude to Bjarne Gram Hansen and Ida Hilden for enabling and guiding us in exploring scientific avenues.

## Appendix

**Supplementary Figure 1.**
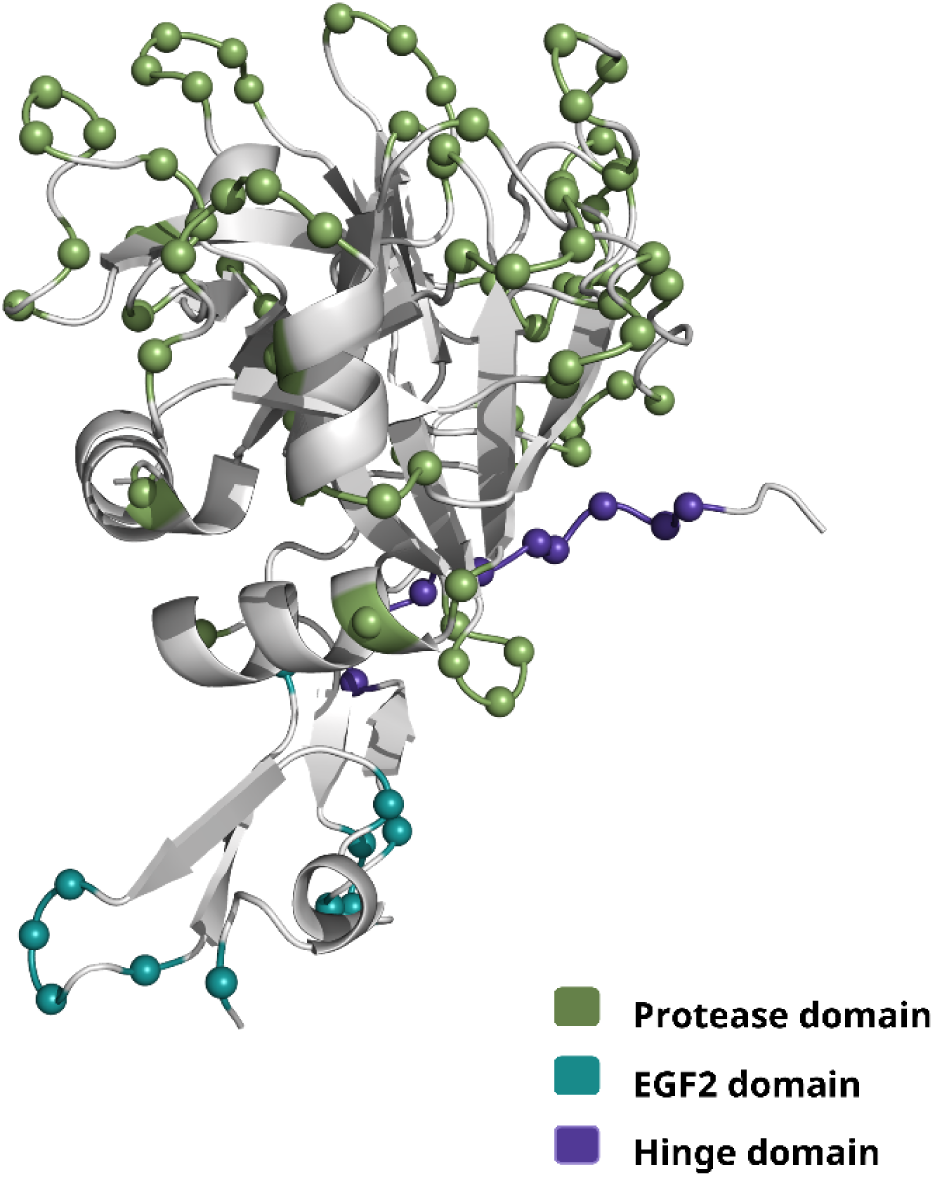
*Distribution of C-alpha positions of N-linked glycans into three domains of fIXa protease domain(smudge green spheres), hinge region (purpleblue spheres), and EGF2 domain (deepteal spheres). Figure was made with PyMOL*(*27*).

### Predict loop structure for matching

The secondary structure was assign using dssp implemented in Rosetta:

rosetta_scripts.linuxgccrelease -database database -parser:protocol dssp.xml -in:file:s PDBFILE @flags

where the xml file, dssp.xml, was given as:

<ROSETTASCRIPTS>

<SCOREFXNS>

<enzdes weights=talaris2013 />

<soft weights=ligand_soft_rep />

</SCOREFXNS>

<TASKOPERATIONS>

<InitializeFromCommandline name=init/>

<SetIGType name=linmem_ig lin_mem_ig=1/>

<LimitAromaChi2 name=limchi2/>

</TASKOPERATIONS>

<FILTERS>

</FILTERS>

<MOVERS>

\# Minimization of complex - no design allowed

<TaskAwareMinMover name=min bb=0 chi=1 jump=1 scorefxn=enzdes task_operations=init/> # Packing of rotamers making sure no aromatic with chi2 of 90 degrees

<PackRotamersMover name=repack scorefxn=enzdes task_operations=init,limchi2/>

<ParsedProtocol name=min_repack_min>

<Add mover=min/>

<Add mover=repack/>

<Add mover=min/>

</PARSEDPROTOCOL>

<Dssp name=dssp reduced_IG_as_L=0/>

</MOVERS>

<PROTOCOLS>

<Add mover_name=dssp/>

</PROTOCOLS>

</ROSETTASCRIPTS>

using the following flags:

-ignore_zero_occupancy false

-packing

-use_input_sc

-ex1

-ex2

-ex1aro

-ex2aro

-linmem_ig 10

-no_optH false

-flip_HNQ

-no_his_his_pairE

-nblist_autoupdate

### Match for N-glycan using Rosetta Matcher

The geometrical search is performed using the executable below match.linuxgccrelease -in:file:s PDBFILE -match:lig_name NAG - match:geometric_constraint_file CSTFILE extra_res_fa NAG.fa.params -match:scaffold_active_site_residues POSITIONFILE where the geometrical restraints have been specified for bond and angle:

CST::BEGIN

TEMPLATE:: ATOM_MAP: 1 atom_name: C1 O3 C2 TEMPLATE:: ATOM_MAP: 1 residue3: NAG TEMPLATE:: ATOM_MAP: 2 atom_type: NH2O, TEMPLATE:: ATOM_MAP: 2 residue1: N CONSTRAINT:: distanceAB: 1.40 0.10 1000.00 1 0

CONSTRAINT:: angle_A: 105.00 1.00 100.00 360.00 1

CONSTRAINT:: angle_B: 120.00 5.00 100.00 360.00 1

CONSTRAINT:: torsion_A: -164.20 10.00 50.00 360.00 1

CONSTRAINT:: torsion_B: -120.00 10.00 50.00 360.00 1

CONSTRAINT:: torsion_AB: -117.00 360.00 0.00 360.00 12 CST::END

The following positions were screened:

1 3 4 7 8 9 10 11 12 13 14 20 21 22 23 24 25 33 34 39 43 44 45 46 47 48 49 54 55 56 58 59 60

61 62

63 64 65 76 77 78 79 81 82 83 85 86 87 88 89 90 96 97 98 99 100 101 103 104 106 107 108

110 111

112 123 130 131 133 134 135 136 137 138 140 141 143 151 152 160 161 162 163 164 165 166

167

172 173 174 175 176 177 178 180 181 182 183 184 185 186 187 193 194 195 196 207 208 209

210

211 212 213 214 215 224 233 234 235 236 237 238 239 240 246 247 248 253 254 255 256 257

262

263 264 265 269 270 271 272 273 274 278 279 280 281 282 283 284 285 286 287 288 289 290

291

292 293

### Geometrical restraint score for each match

To evaluate each match we score how well the geometrical parameters are obtained on the structure rosetta_scripts.linuxgccrelease -database rosetta_database -parser:protocol score_pose.xml - in:file:s PDBFILE -extra_res_fa NAG.fa.params @flags -parser_read_cloud_pdb 1 the flag file is defined the following way:

-preserve_header 1

-parser_read_cloud_pdb 1

-packing

-use_input_sc

-ex1

-ex2

-ex1aro

-ex2aro

-linmem_ig 10

-no_optH false

-flip_HNQ

-no_his_his_pairE

-nblist_autoupdate

-ignore_zero_occupancy false and score_pose.xml is:

<ROSETTASCRIPTS>

<TASKOPERATIONS>

<InitializeFromCommandline name=init/>

<ReadResfile name=res filename=resfile />

<LimitAromaChi2 name=limchi2/>

</TASKOPERATIONS>

<SCOREFXNS>

<enzdes weights=talaris2013_cst />

</SCOREFXNS>

<MOVERS>

<AddOrRemoveMatchCsts name=addcst cstfile=“nglycan.cst” cst_instruction=add_new/> # Minimization of complex - no design allowed

<TaskAwareMinMover name=min bb=0 chi=1 jump=1 scorefxn=enzdes task_operations=init/> # Packing of rotamers making sure no aromatic with chi2 of 90 degrees

<PackRotamersMover name=repack scorefxn=enzdes task_operations=init,limchi2,res/>

<ParsedProtocol name=min_repack_min>

<Add mover=min/>

<Add mover=repack/>

<Add mover=min/>

</PARSEDPROTOCOL>

</MOVERS>

<FILTERS>

<EnzScore name=cst_score scorefxn=enzdes whole_pose=1 score_type=cstE energy_cutoff=10.0/>

</FILTERS>

<PROTOCOLS>

<Add mover=addcst/>

<Add mover=min_repack_min/>

<Add filter=cst_score/>

</PROTOCOLS>

<OUTPUT scorefxn=enzdes/>

</ROSETTASCRIPTS>

### Python script to convert dssp output to position file

To sample loop regions of the protein, dssp is used to determine loop regions in the protein. The dssp

output from Rosetta is converted into position file used by the match algorithm. # Convert the DSSP output from Rosetta into a

\# position file used for the match algorithm #

\# python convert_dssp_to_positionfile.py dsspfile #

\# generates the position file # position.pos

import sys

inputfile = sys.argv[1] tmp_pos = []

with open( inputfile ,’r’ ) as f:

for line in f:

tmp_pos.append( line )

with open(“position.pos”,’w’) as f:

for dssp in range(len (tmp_pos[0]) ):

if tmp_pos[0][dssp] == ‘L’ :

f.write( str(dssp+1) +“L”)

### Calculating DSSP of crystal structure (PDB ID: 1RFN)

The dssp is computated using Rosetta to assign secondary structure LELLEELLLLLLLLEEEEELLLLLLEEEEEEELLEEEELHHHLLLLLLLEEEELLLELLLLLLLLEE EEEEEEEELLLLELLLELLLLLLEEEEELLLLLLELLELLLELLLHHHHHHHHHHLEEEEEELLEL

LLLLLELLELEEEEEEELLHHHHHHHLLLLLLLLEEEELLLLLLLELLLLLLLLEEEEELLLLEEE EEEEEEELLLLLLLLLEEEEEHHHLHHHHHHHHLLLLLLLLHHHHHLLLEEEELLLLLEEEELLL LEEELLLLLLEEELLLLLLLLLLLLLLLL

### Characterization of N-linked variants

**Supplementary Table 2.**
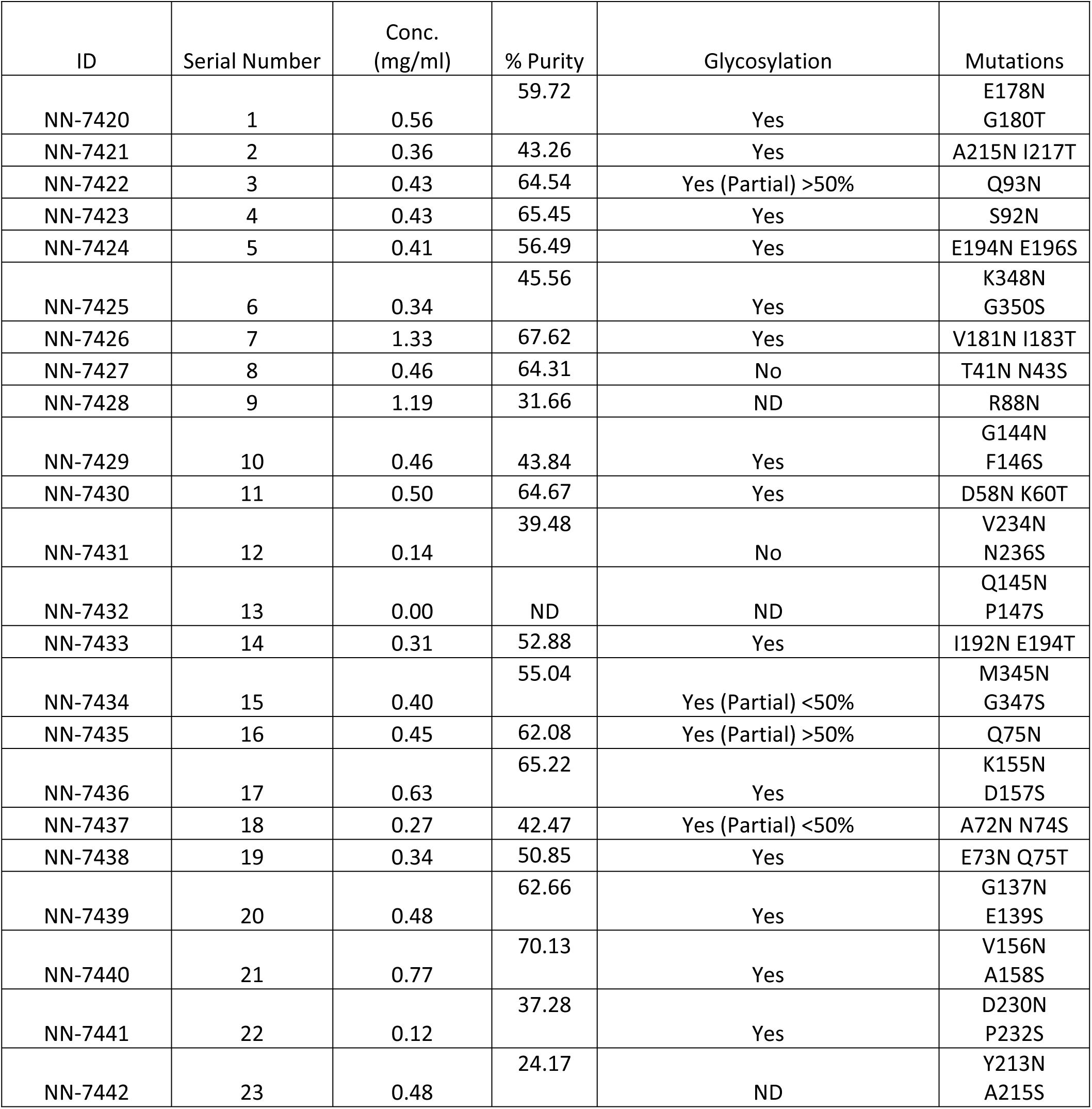
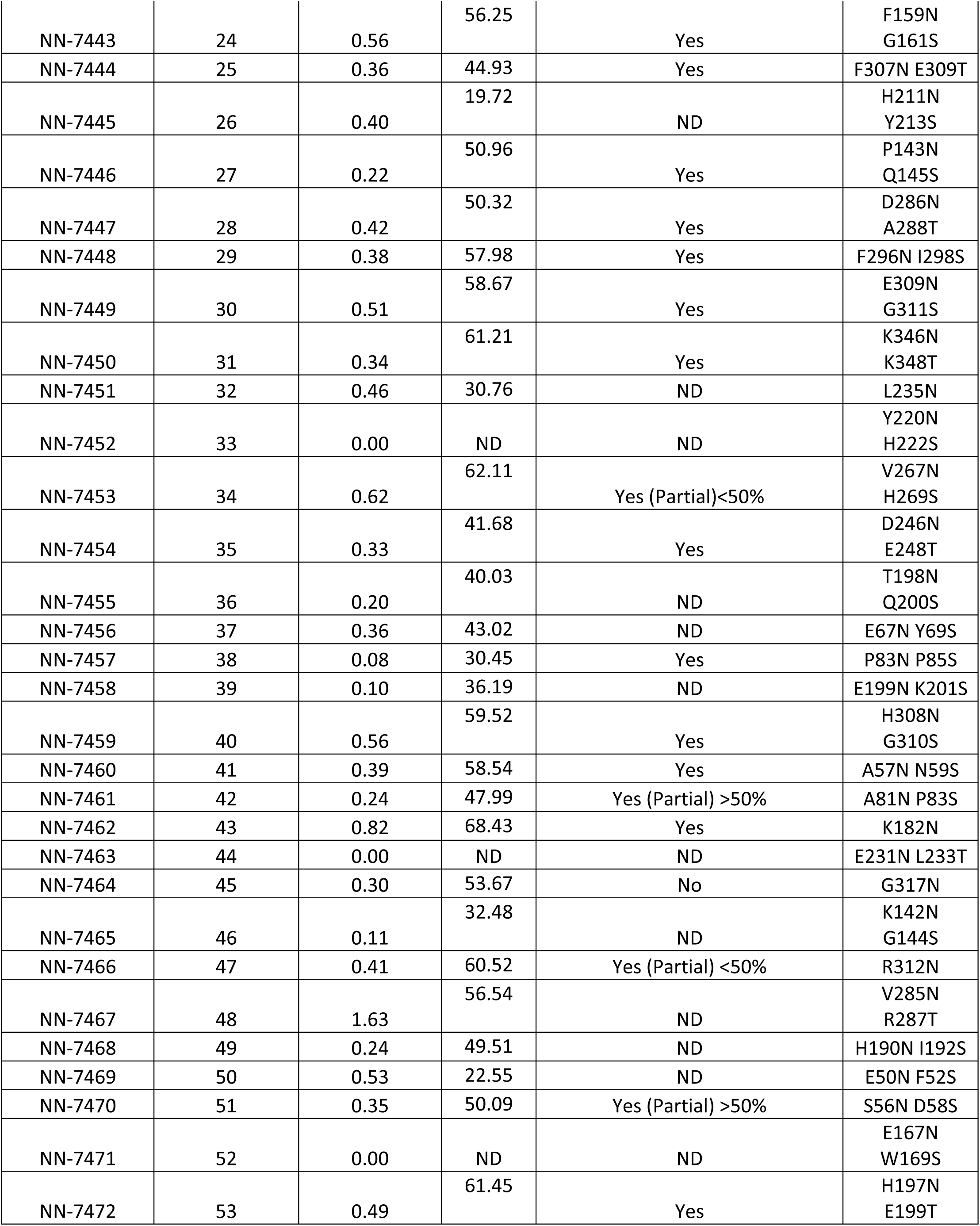
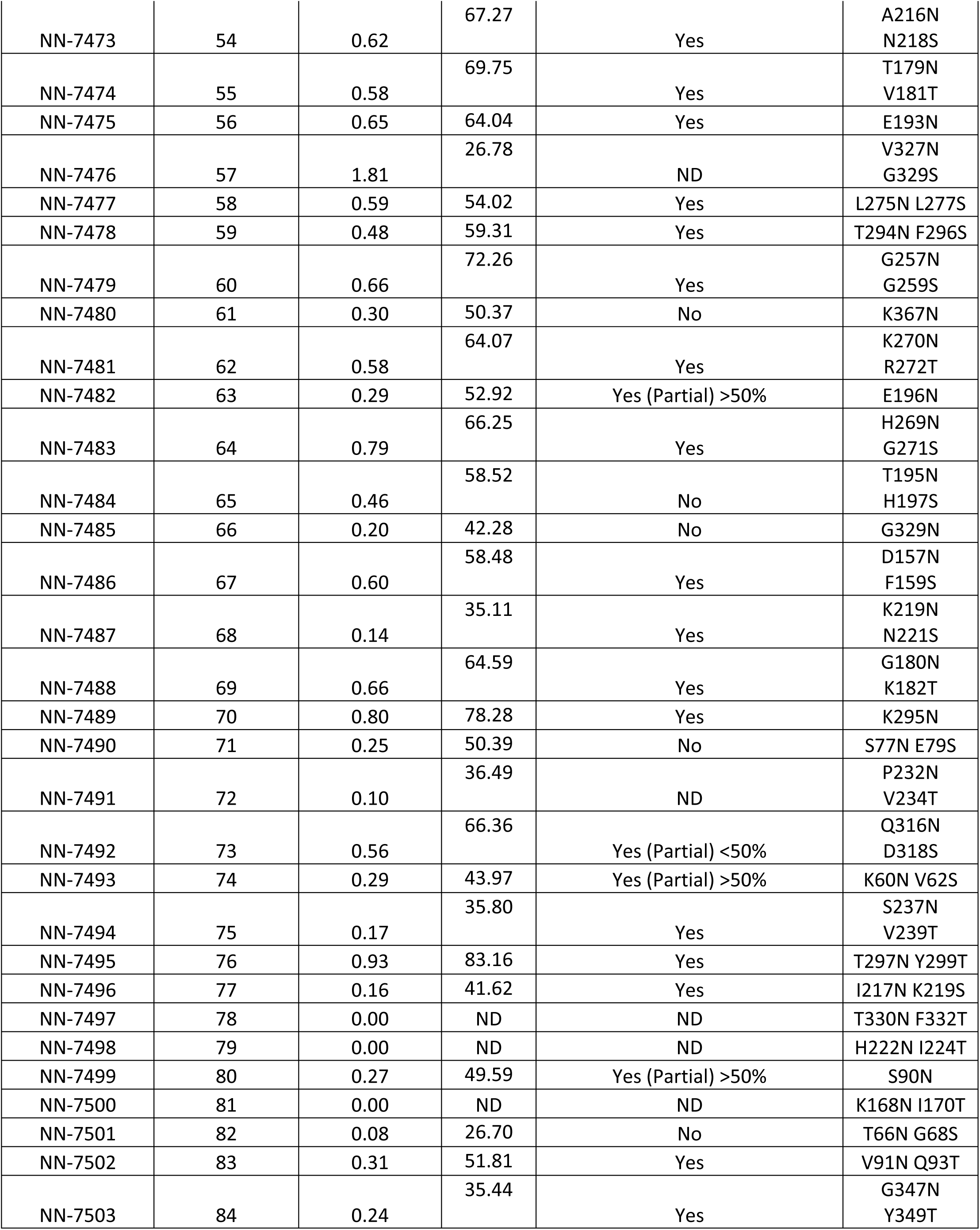
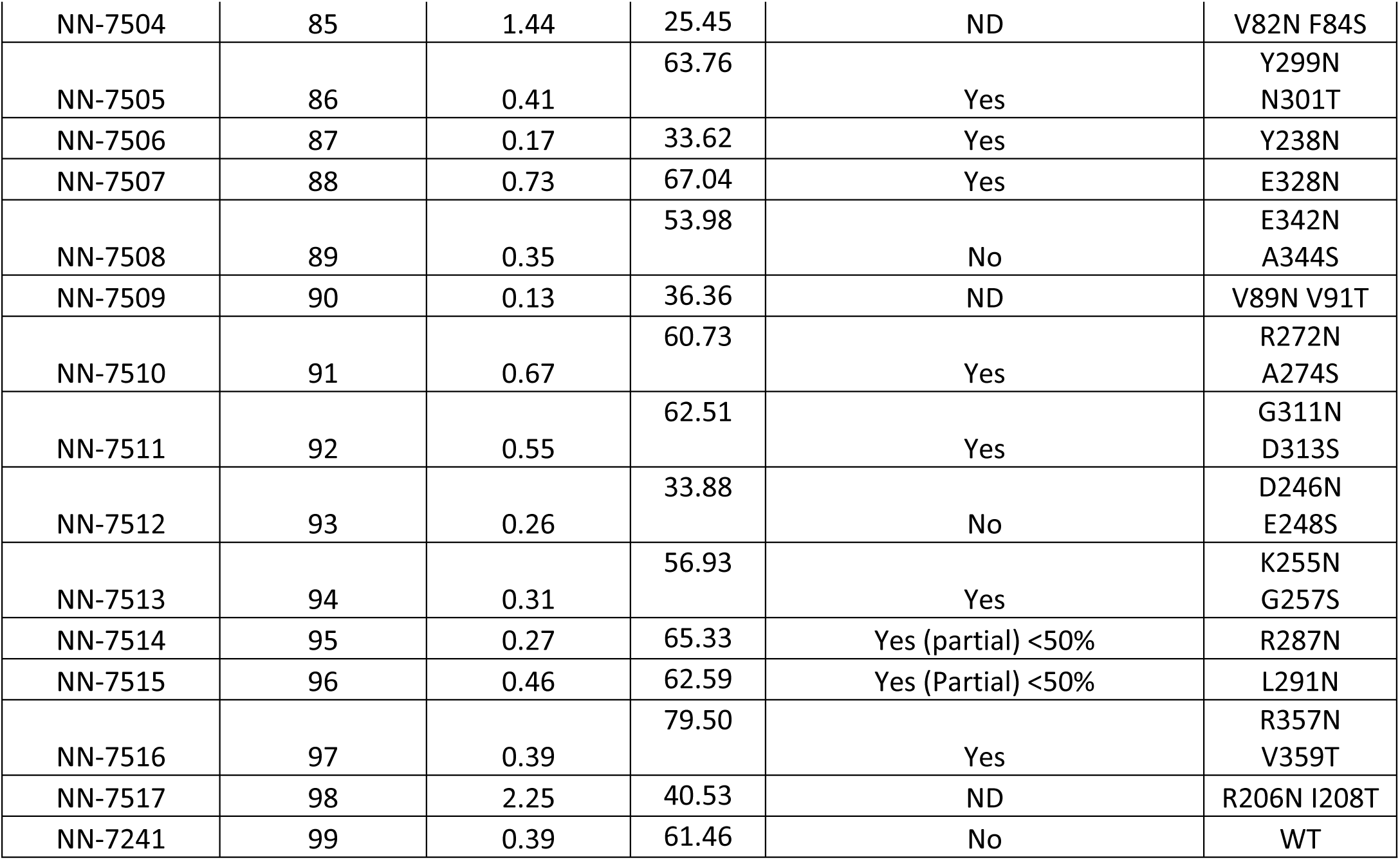
*Engineered N-linked glycosylated variant with ID, Serial Number, concentration, purity, assessment of glycosylation, and mutations*.

**Supplementary Figure 2.**
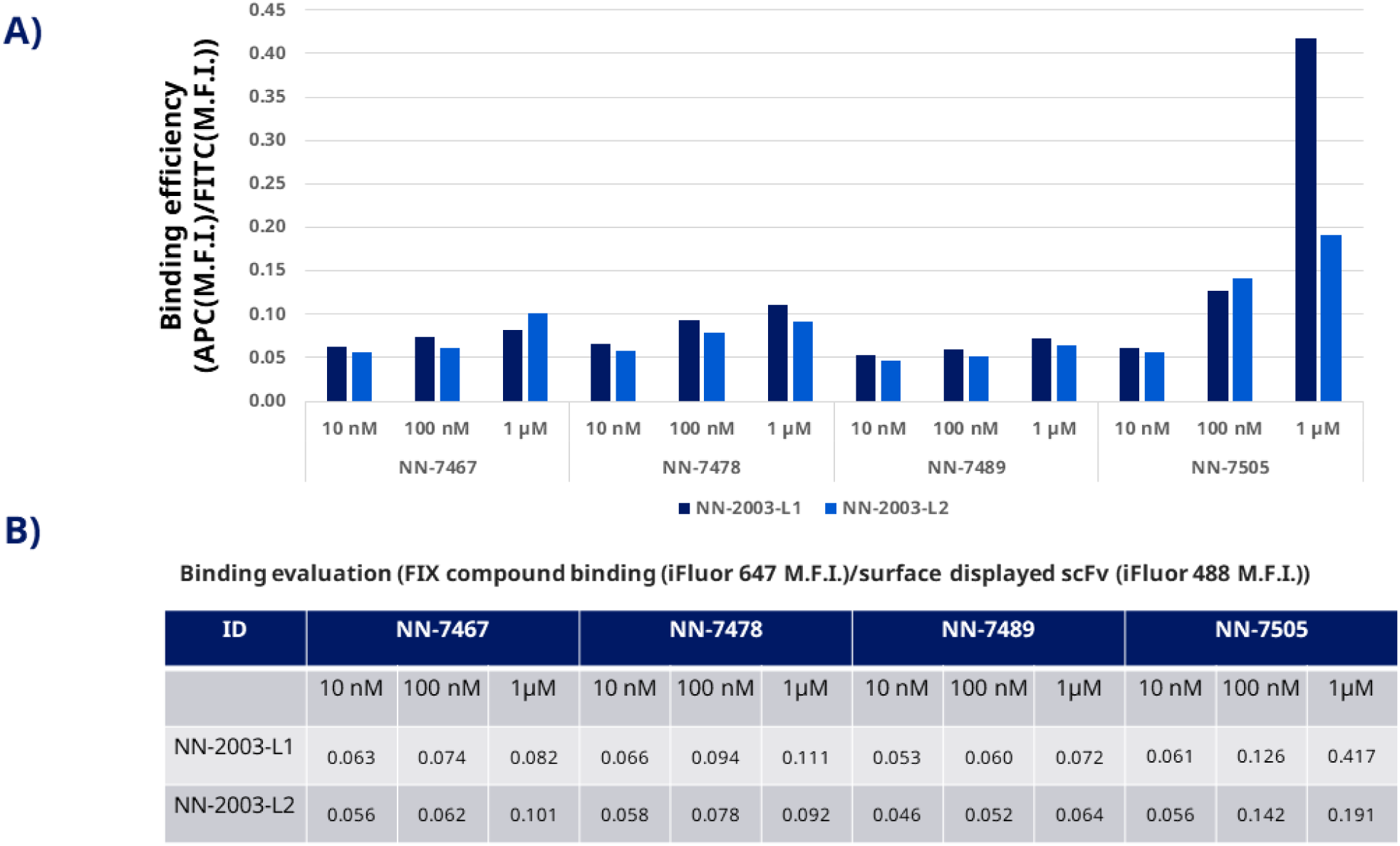
*Binding study of NN-2003-L1 and NN-2003-L2 against four N-linked glycosylated fIXa variants using yeast surface display. A) shows the binding efficiency against the four variants. B) Tabular data of the binding study*.

### Engineered N-linked glycosylated sequences of fIXa

**Supplementary Table 3.**
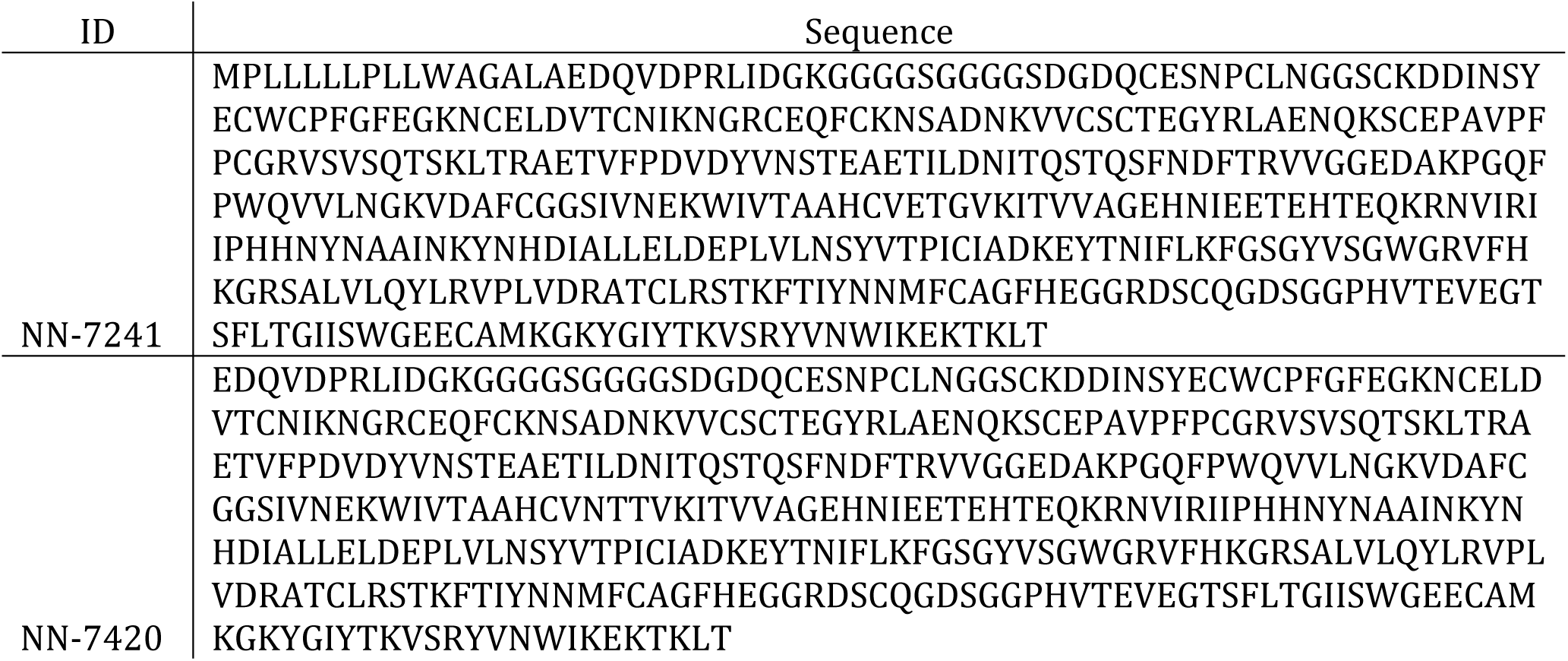
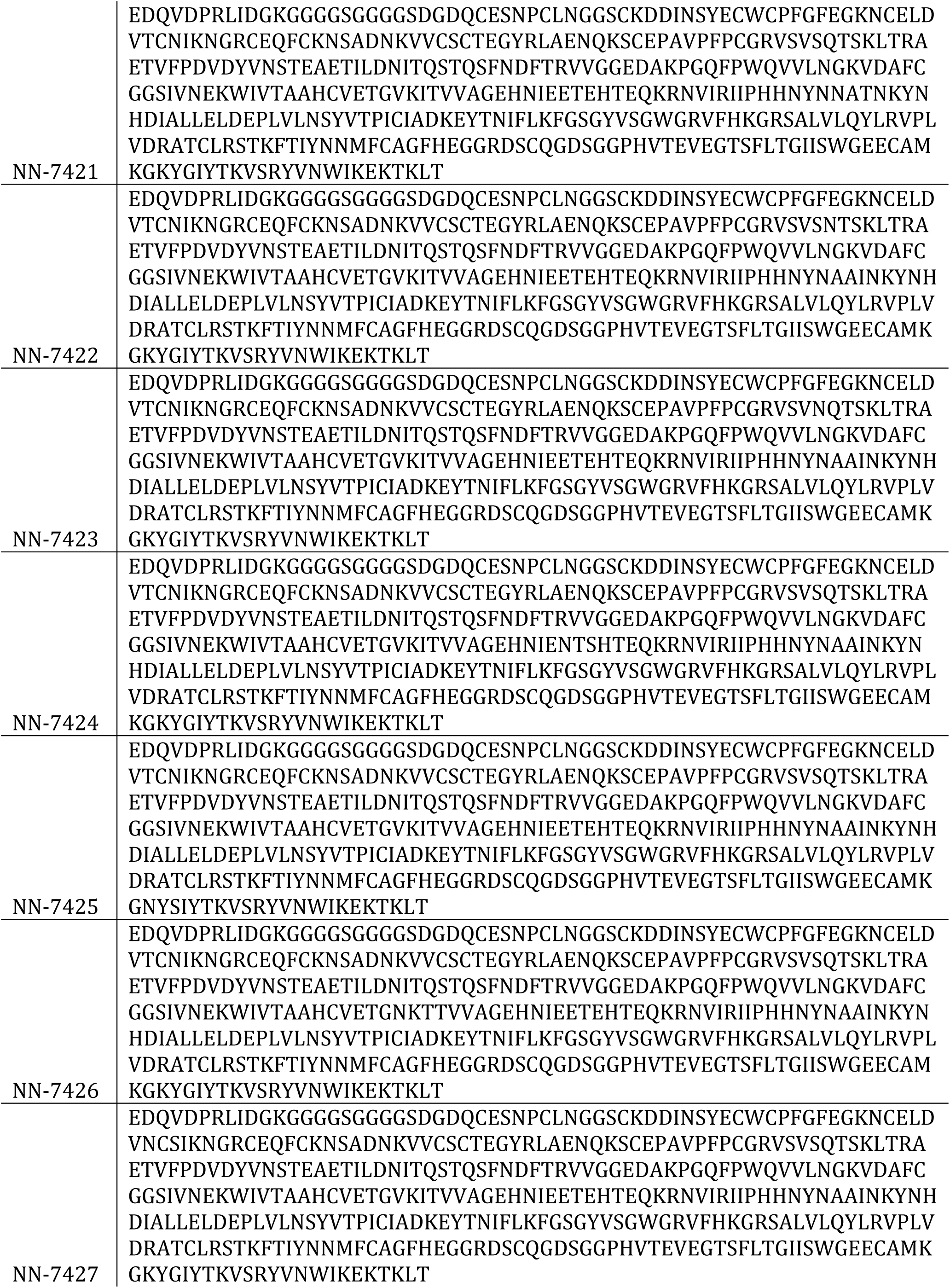
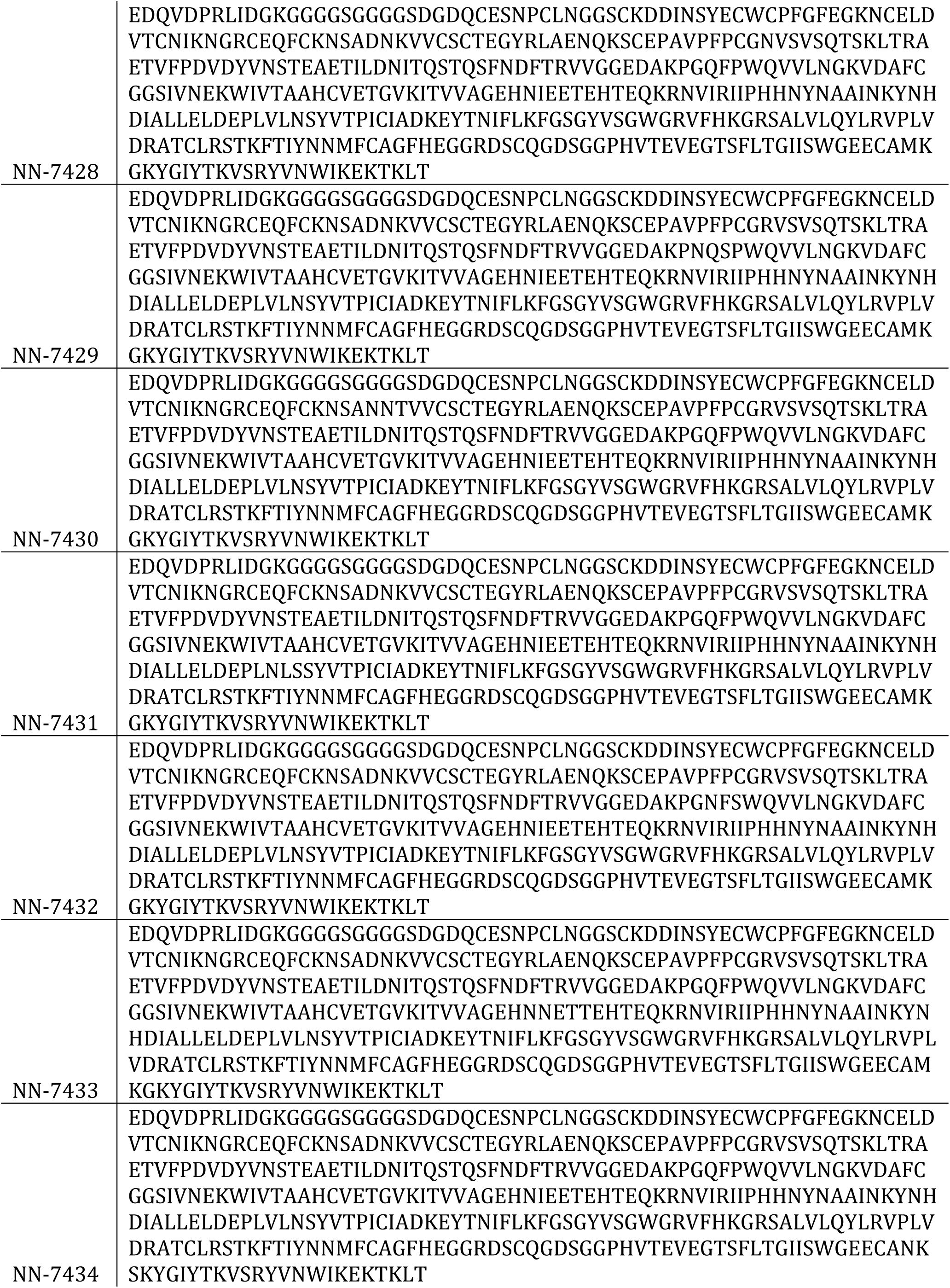
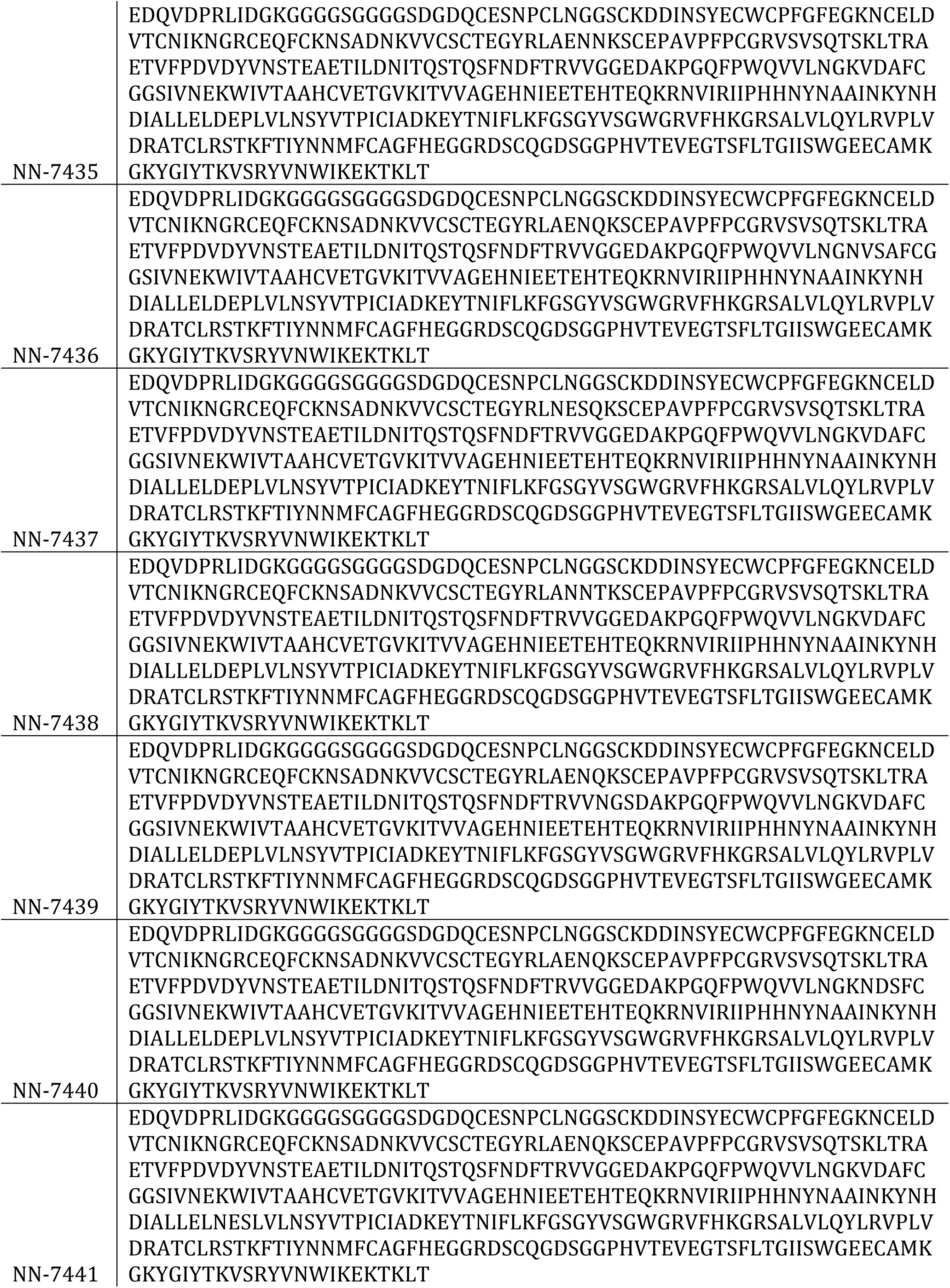
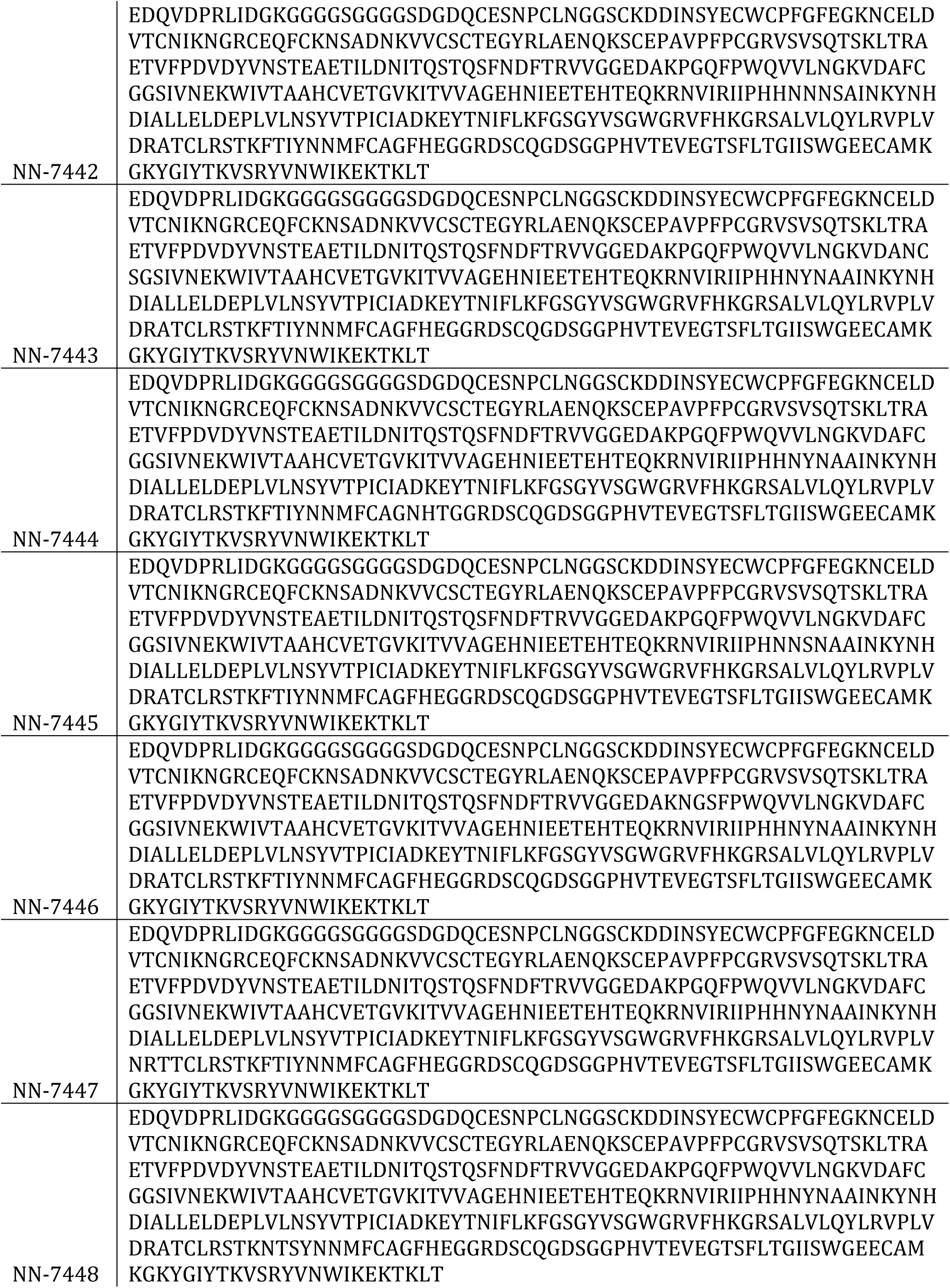
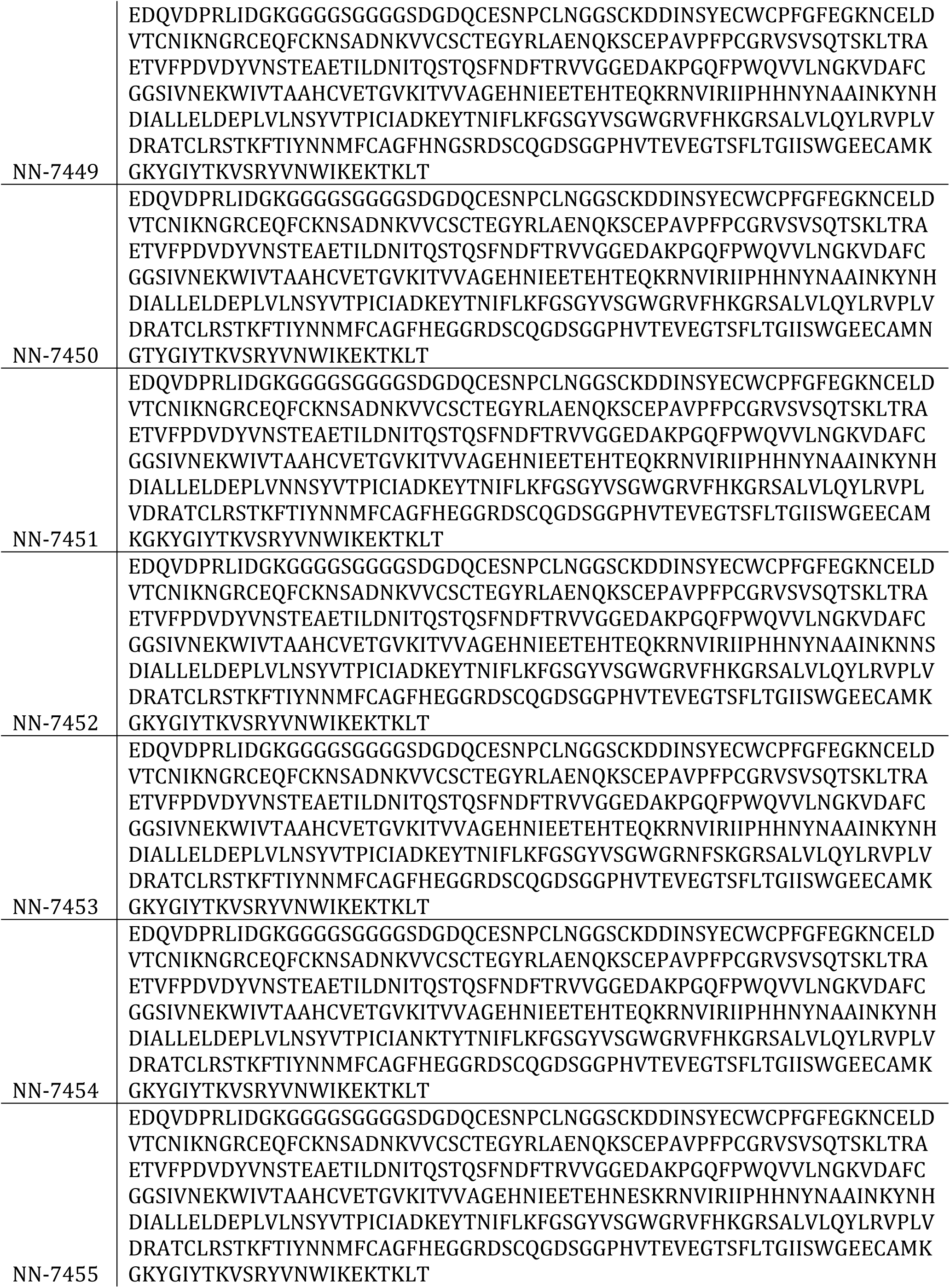
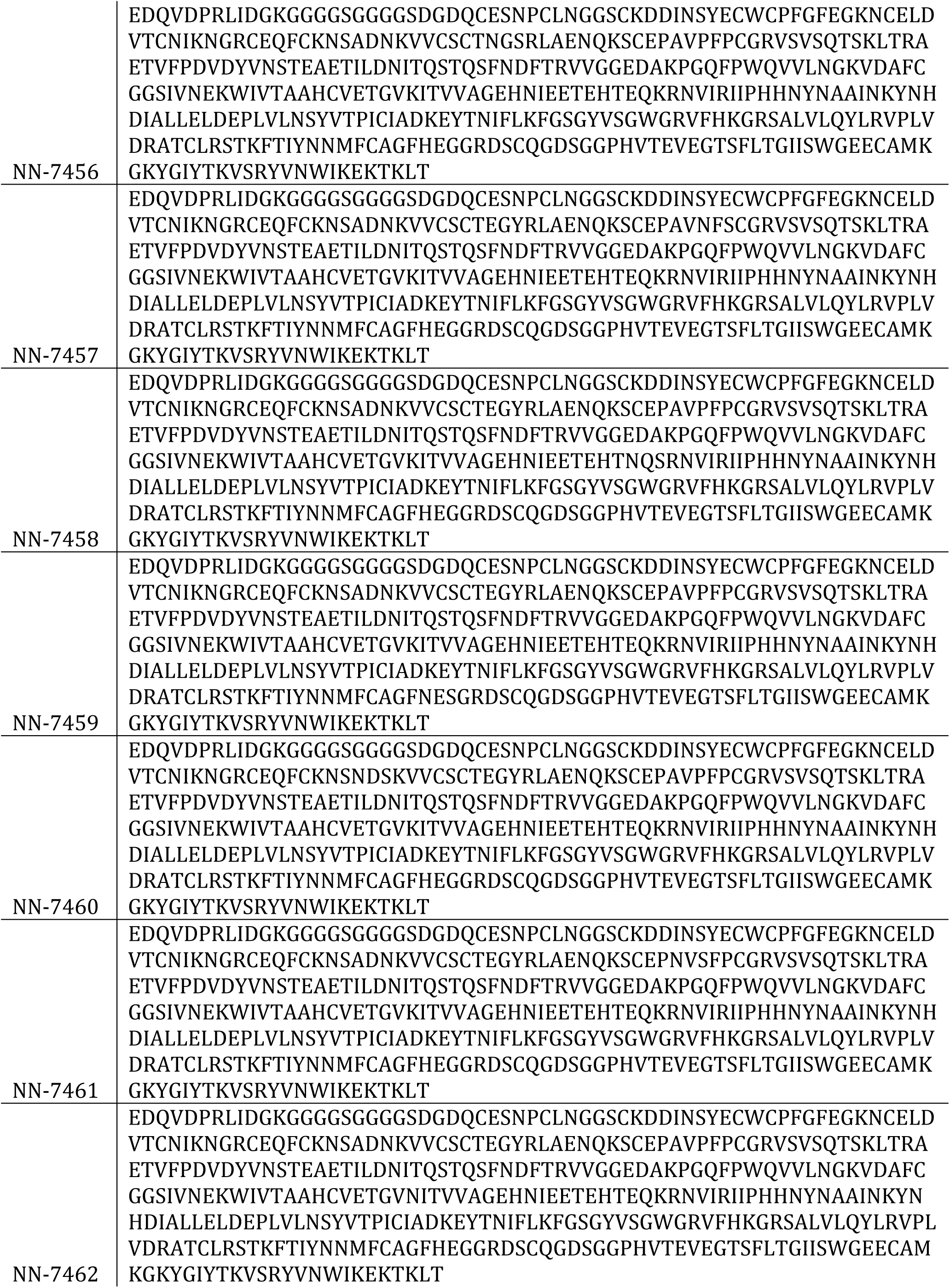
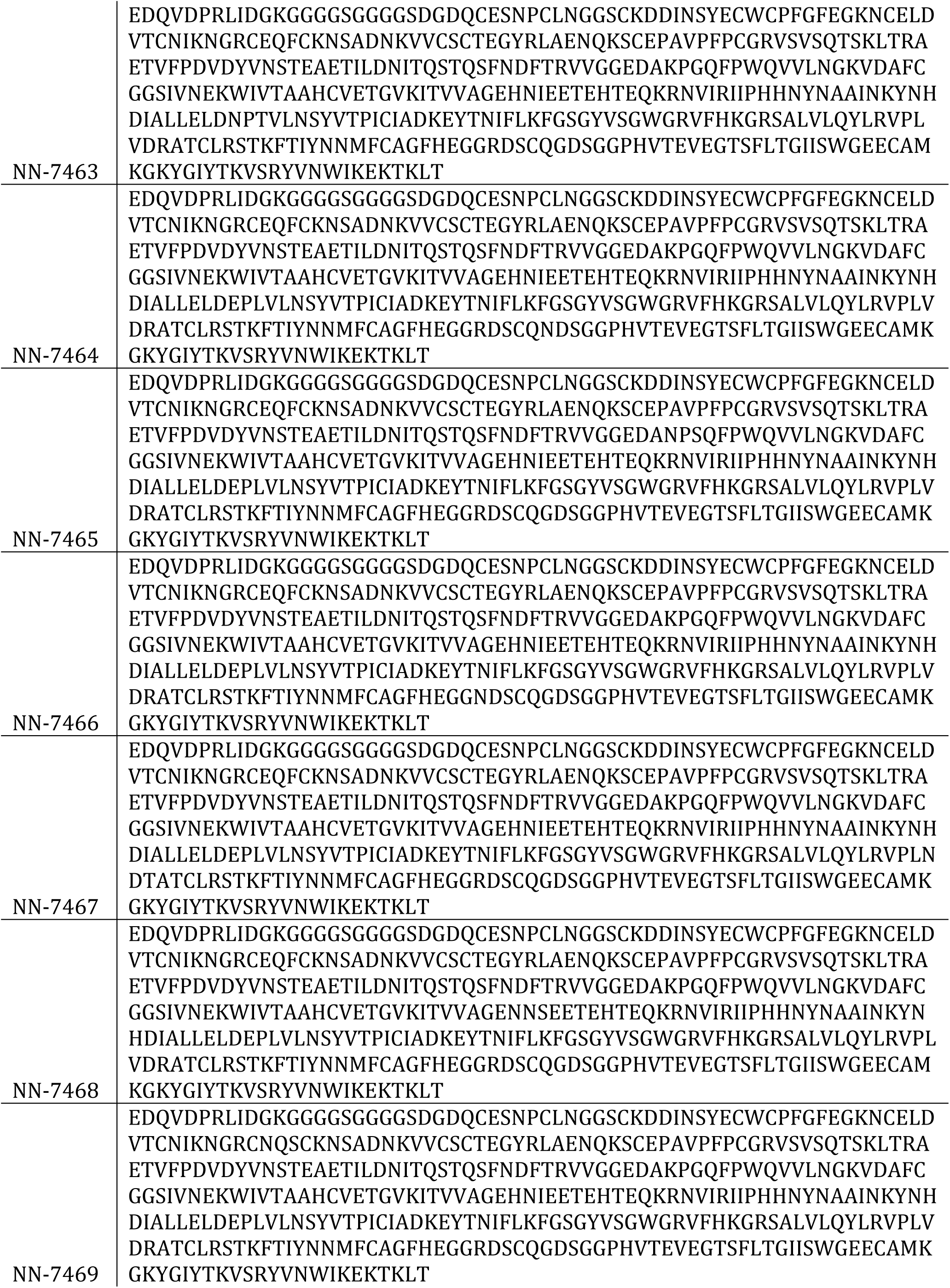
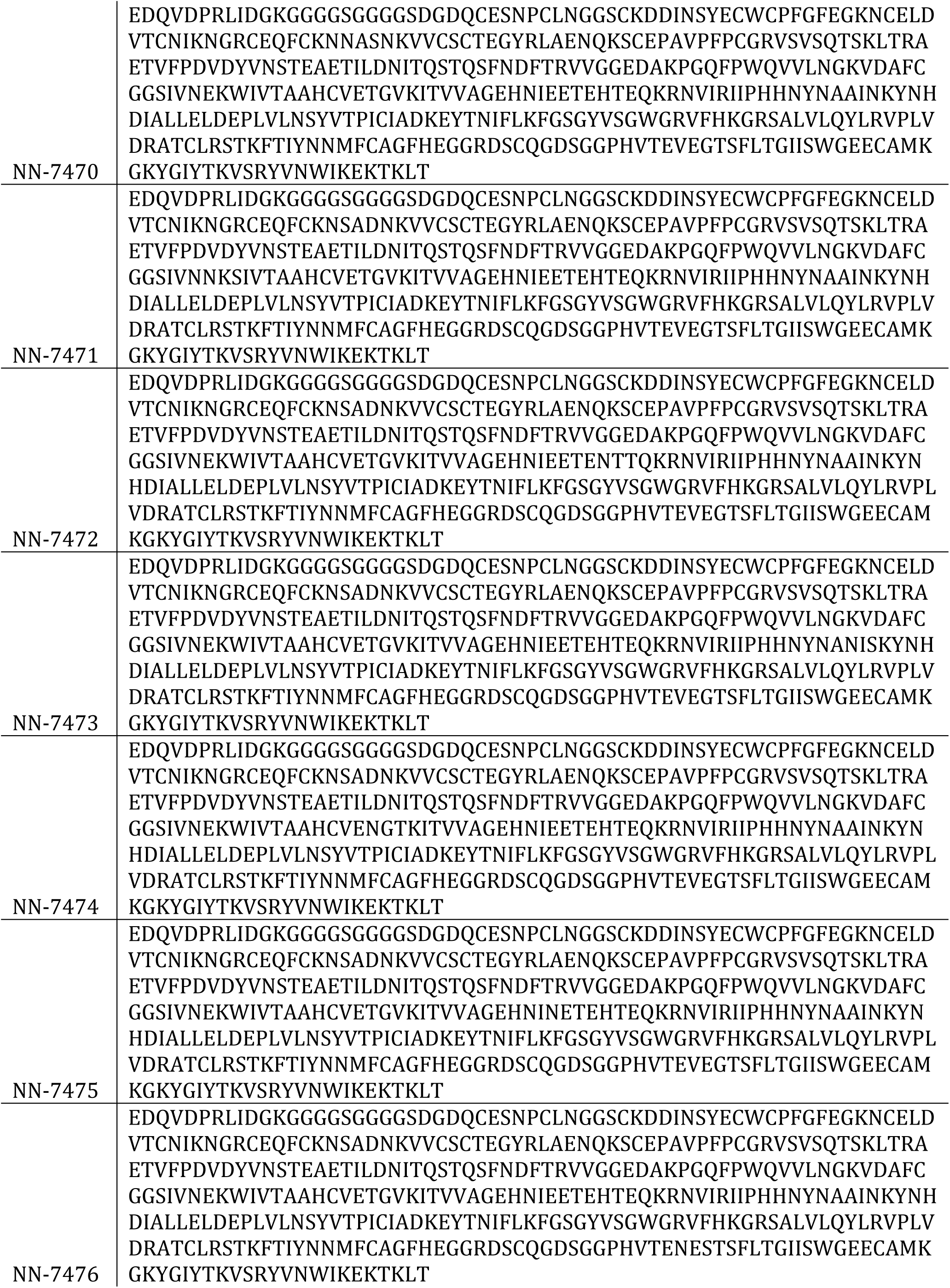
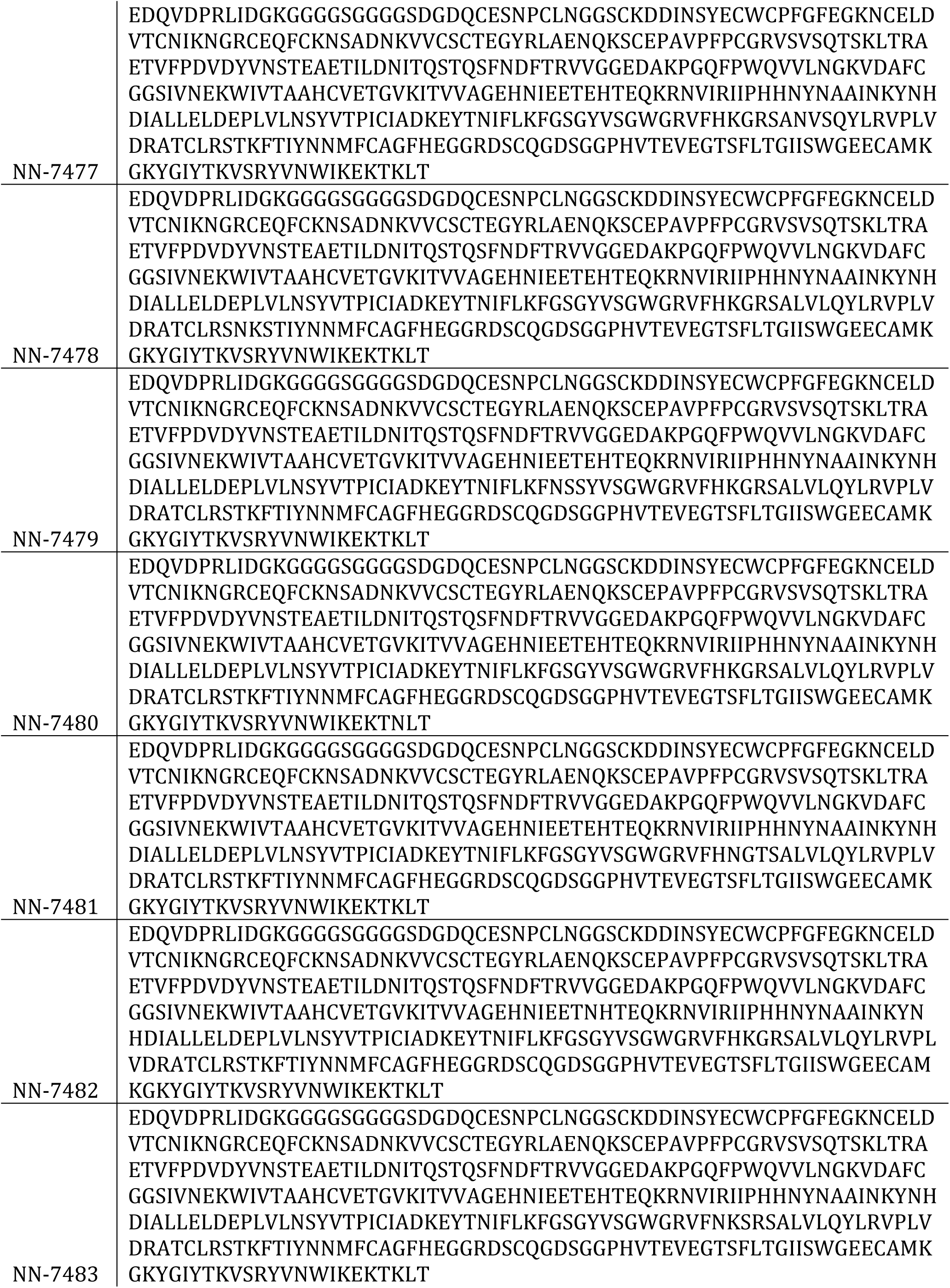
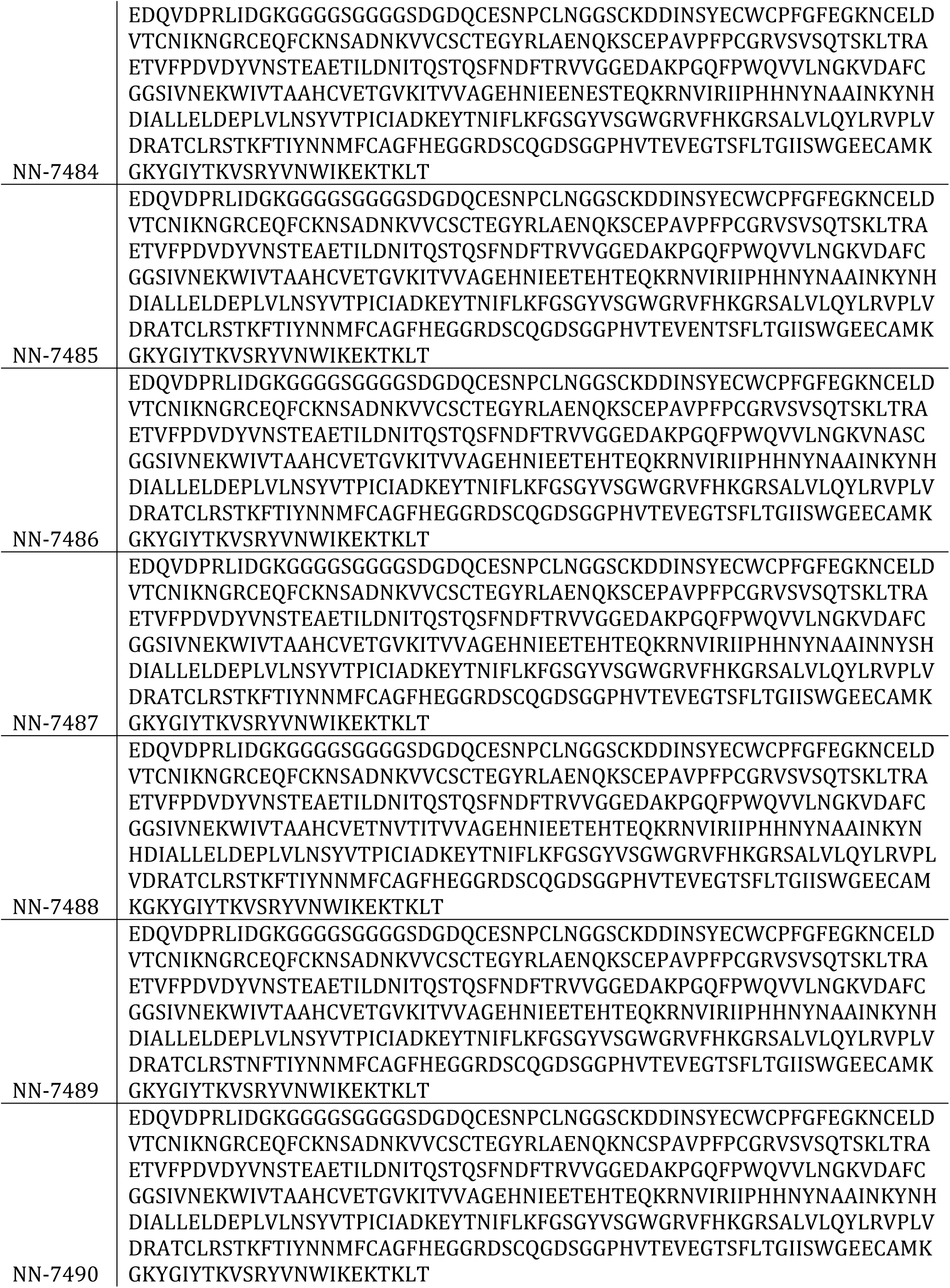
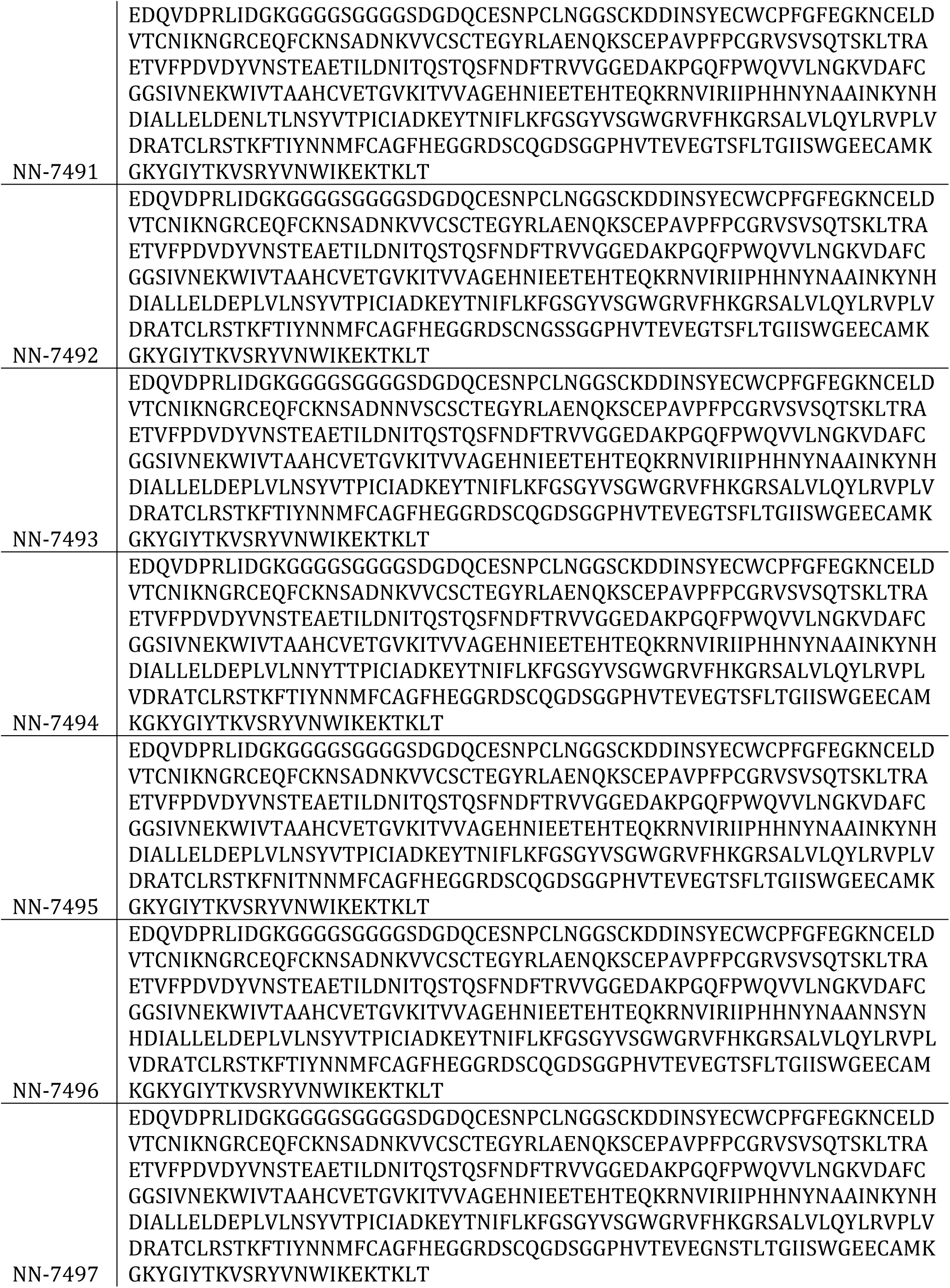
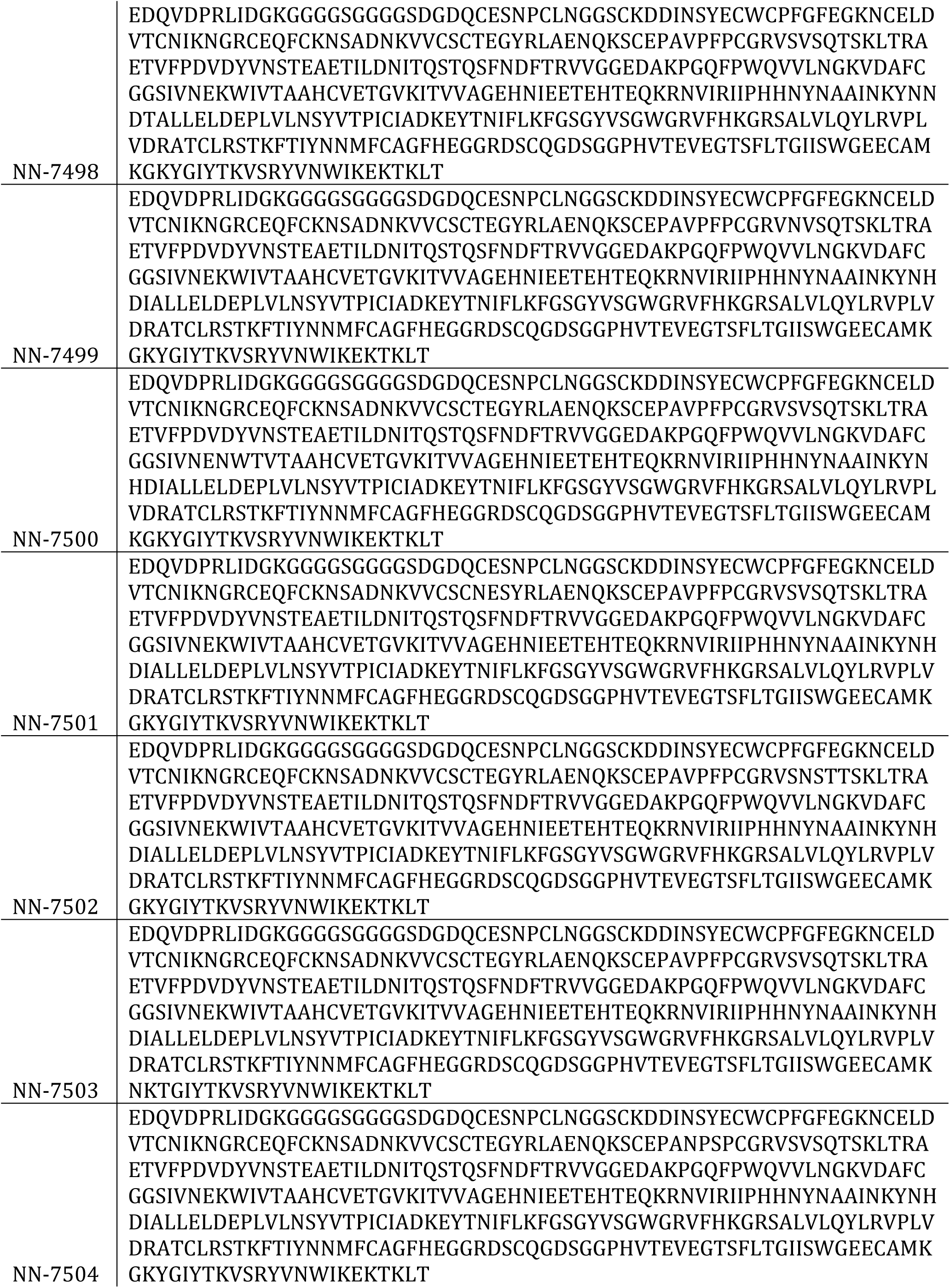
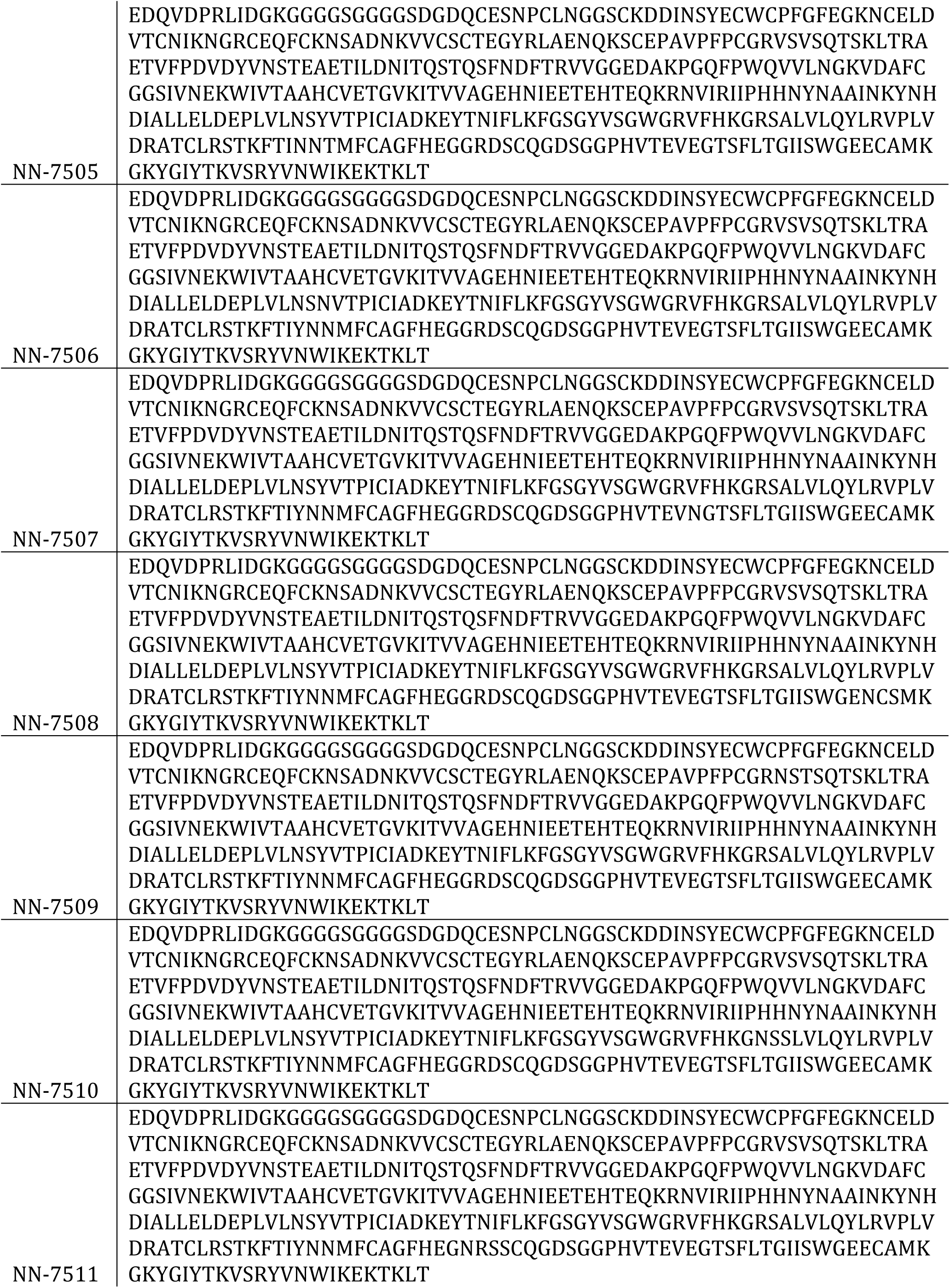
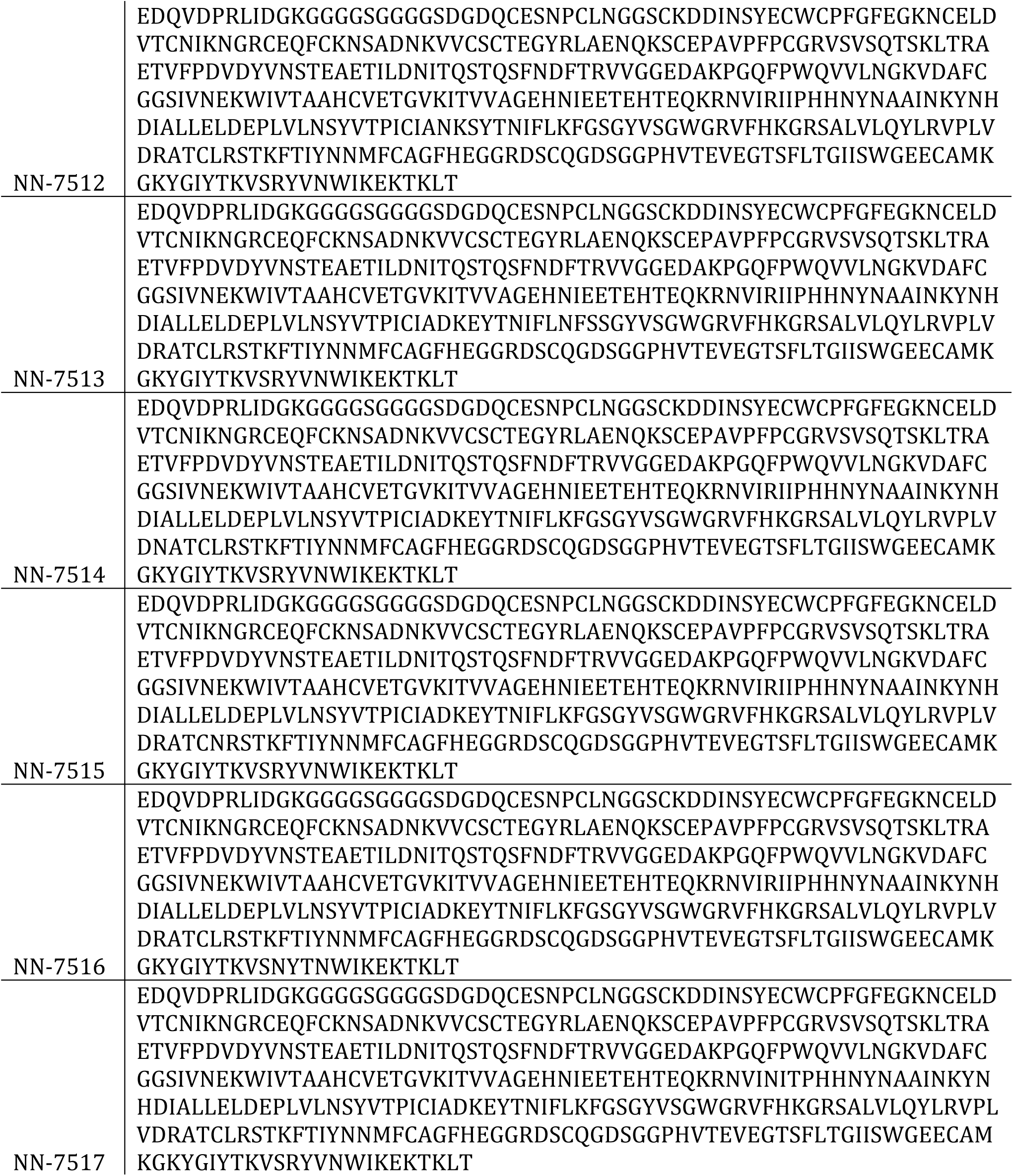
*Sequences of N-linked glycosylated variants*

